# Dual Dean entrainment with volume ratio modulation for efficient co-encapsulation: Extreme single-cell indexing

**DOI:** 10.1101/2021.04.08.439026

**Authors:** Jack Harrington, Luis Blay Esteban, Jonathan Butement, Andres F. Vallejo, Simon I. R. Lane, Bhavwanti Sheth, Maaike S. A. Jongen, Rachel Parker, Patrick S. Stumpf, Rosanna C. G. Smith, Ben D. MacArthur, Matthew J. J. Rose-Zerilli, Marta E. Polak, Tim Underwood, Jonathan West

## Abstract

The future of single cell diversity screens involves ever-larger sample sizes, dictating the need for higher throughput methods with low analytical noise to accurately describe the nature of the cellular system. Current approaches are limited by the Poisson statistic, requiring dilute cell suspensions and associated losses in throughput. In this contribution, we apply Dean entrainment to both cell and bead inputs, defining different volume packets to effect efficient co-encapsulation. Volume ratio scaling was explored to identify optimal conditions. This enabled the co-encapsulation of single cells with reporter beads at rates of ~1 million cells/hour, while increasing assay signal-to-noise with cell multiplet rates of ~2.5% and capturing ~70% of cells. The method, called Pirouette-seq, extends our capacity to investigate biological systems.

**TOC Abstract:** Pirouette-seq involves cell and reporter bead inertial ordering for efficient co-encapsulation, achieving a throughput of 1 million cells/hour, a 2.5% multiplet rate and a 70% cell capture efficiency.

## Introduction

Elucidating the origins, development and fate of cellular systems is at the forefront of biological enquiry. Increasingly single cell next generation sequencing (NGS) profiling is used to provide a comprehensive map of the cellular population linked to each cell’s underlying processes and their role in system biology. In essence, these experiments involve compartmentalising single cells with reporter beads to capture and encode a cell’s biological properties prior to delivery to a NGS and bioinformatics pipeline. Cell and bead co-encapsulation requires small volume liquid handling, a central strength of microfluidics^1^. First, elastomeric Quake valves^2^ were used to compartmentalise single cells within an addressable array, marking the beginnings of single-cell diversity screens^3, 4^. Dramatic increases in throughput (*e.g*. 10,000’s of cells) emerged from using nanowell arrays^5–7^ and droplet microfluidics^8, 9^.

Following these pivotal technology developments, single-cell analysis is gaining pace with the biology community aiming to decipher ever-larger cellular systems. Cell Atlas reference maps, CRISPR and compound library single-cell screening projects typify this trend, with the scalability of the continuous flow droplet microfluidics format suitable for matching such large experiments^10^. However, efforts in this direction face a fundamental problem: Cells are randomly encapsulated. The probability of cells being encapsulated in a droplet is described by the Poisson statistic;

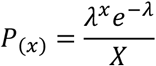

where *P*_(*X*)_ is the probability of *X* number of cells being packaged in a droplet and *λ* the mean number of cells per droplet. Dilute cell suspensions (*λ* <0.1) are used to reduce the probability of co-encapsulating multiple cells and prevent their biology being scrambled during barcoding. As a consequence throughput is greatly limited and cell multiplets cannot be completely excluded, resulting in a trade-off between throughput and analytical noise: For most experiments a signal-to-noise of >20 (<5% multiplet rate) is acceptable which drastically limits cell concentrations. To compound the problem, solid reporter beads are also delivered as dilute suspensions to avoid clogging. This collides two Poisson statistics as a joint probability distribution (JPD) in which cell and bead coupling rates are necessarily low (<1% of droplets, see **Supplementary Information, Figure 1**). Alternatively, hydrogel beads can be delivered to most droplets (50–95%) using packed flows. This reduces the problem to a single Poisson statistic allowing the majority of cells to be captured, albeit necessitating lower droplet generation rates to produce equivalent throughput to solid bead systems.

**Figure 1.**
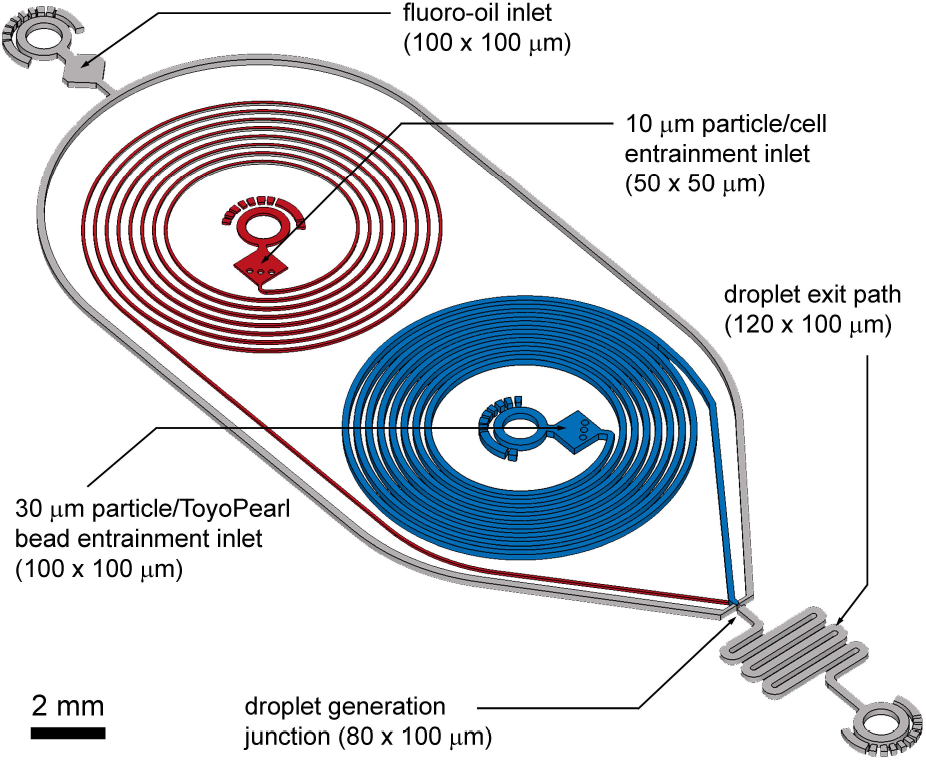
Pirouette-Seq microfluidics circuit for dual Dean entrainment and droplet generation (to scale). Two-layer fabrication is used to effect entrainment of the dissimilarly-sized reporter beads and cells; spiral channels with a height of 50 μm were used for cell entrainment, and channels 100 μm in height were used for bead entrainment and droplet generation.

Inertial microfluidic formats offer the enticing possibility of the periodic delivery of cell and beads into droplets to free assays from the limitations of the Poisson statistic. During the high velocity transport (*Re* > 1) of particle-laden flows where particle diameters approach microchannel dimensions (*a/Dh* > 0.07) particle interaction with the underlying flow field can be predicted by the Reynolds particles number, *Re_p_* = *Re(a/Dh)^2^* ≥ 0.1^11–13^. In this regime the parabolic velocity profile introduces an appreciable shear-gradient lift force that becomes countered by the wall effect lift force to produce an equilibrium position, focusing the particles within the same streamlines. This has the effect to increase the local particle concentration resulting in particle trains^14^ with the interplay between viscous disturbance and inertial lift forces producing an equilibrium defining the inter-particle spacing^15^. Using straight channels these principles have been applied to the formation of *‘microfluidic crystals’*^16^ and deterministic cell encapsulation in droplets^17, 18^. Introducing secondary Dean flows (*De = Re(D_h_/R)^1/2^*, where *R* is the channel radius of curvature) created by high velocity transport in curved channels increases migration to attain faster particle focusing and train formation^19, 20^. With the benefit of curvature, spiral channels have been used to increase solid bead droplet loading to enhance cell capture efficiencies from ~5% to 20%^21^. This also introduces gains in throughput, but remains limited to dilute cell suspensions dictated by the Poisson statistic. Entraining both beads and cells has proved challenging, again requiring dilute cell suspensions to reduce multiplets^22^.

Reporter beads and cells have dissimilar sizes (ø30 **μ**m v. ø10–15 **μ**m). We reasoned that each requires tailored microfluidic conditions for effective entrainment. In this study, we have developed a two-layer prototype for the effective Dean entrainment of the solid beads and cells (**Figure 1**). The volumetric ratio between beads and cells was investigated to identify operating windows for highly efficient co-encapsulation, surpassing the state of the art: We call the approach Pirouette-seq, a technique enabling the large-scale expansion of single-cell experiments.

## Materials and Methods

### Design

Following Dean entrainment principles we designed a two-layer microfluidic prototype that is illustrated in **Figure 1** and provided as a CAD file (**SI CAD**). The prototype incorporates 50 **μ**m wide and high spiral channels for cell entrainment, and 100 **μ**m wide and high spiral channels for ToyoPearl bead entrainment. These channels combine at an 80-**μ**m-wide droplet generation junction adjoining a 120-**μ**m-wide droplet exit channel. Each spiral channel has 6 turns with a radius minimum of 1.6 mm and maximum of 3.2 mm to produce an overall length of ~100 mm.

### Fabrication and Assembly

Pirouette-Seq devices were fabricated by standard SU-8 photolithography, followed by replication in PDMS. Inlet and outlet ports were prepared using a 1-mm-diameter biopsy punch (Miltex), and then the device was oxygen plasma bonded (Femto, Deiner) to a glass microscope slide. Surfaces were functionalised by flooding the device with 1% (v/v) trichloro(1*H*,1*H*,2*H*,2*H*-perfluorooctyl)silane (Merck) in HFE-7500™ (3M™). Plug and play interconnection between 25G needles on the syringes and the device inlets was achieved using polythene tubing (Smiths Medical, ID 0.38 mm; OD 1.09 mm).

### Particles, Beads and Cells

Monodisperse 10-**μ**m-diameter polystyrene particles (Merck) were suspended in PBS, and monodisperse 20- and 30-**μ**m-diameter polystyrene particles (Merck) and filtered ≤40-**μ**m ToyoPearl beads (HW-65S, Tosoh Biosciences, unfunctionalized ChemGene beads) were suspended in filtered, modified DropSeq Lysis Buffer^9^ (100 mM Tris, pH 7.5, 0.1% Sarkosyl, 10 mM EDTA). Human THP and HEK293 cells were washed and resuspended in filtered PBS with 1% (w/v) BSA. Particle, bead and cell diameter histograms are provided in the **SI Fig. 2**. Particles, beads and cells were retained in suspension using a vertically orientated syringe with a PTFE-coated samarium cobalt disc magnet rotated at low rpm (≤30 a.u., Multi Stirrus™, V&P Scientific) (**SI Fig. 3**). To avoid particles and beads occluding the channel during high concentration delivery, important instructions are provided in the **SI Appendix I**. This method allows the prolonged delivery of high concentration particle suspensions (*e.g*. 1.5 M/mL ToyoPearl beads for >40 minutes).

### Microfluidics

For the generation of 600 pL (CV<2%) droplets at ~1,800 Hz a QX200 (BioRad) fluoro-oil flow rate of 165 **μ**L/min was used with a total, bead and cell, aqueous flow rate of ~65 **μ**L/min. Flow details for volume ratio scaling are provided in **SI Table 1**. High-speed microscopy (Miro Lab310, Vision Research) was used to image entrainment and droplet encapsulation. Video files were pre-processed in ImageJ or directly analysed using custom MATLAB scripts for measuring inter-particle pitch (**SI Appendix II**) and encapsulation (**Appendix III**). Encapsulation results were verified manually.

### Metric Definitions

The signal-to-noise (*S:N*), multiplet rate (*MR*), throughput (*TP*) and capture efficiency (*CE*) performance metrics describing droplet co-encapsulations are described by:

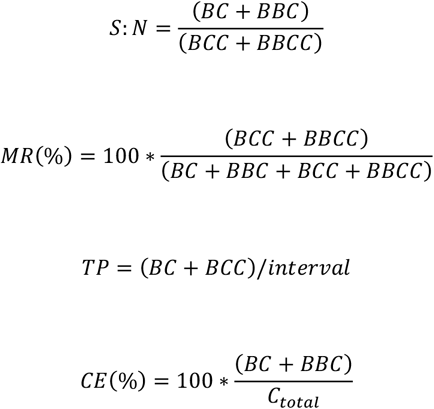

where *B* denotes a 30 **μ**m polystyrene particle or ToyoPearl bead per droplet, *C* denotes a 10 **μ**m polystyrene particle or cell, *BB* denotes 2 or more particles or beads, *CC* denotes 2 or more 10 **μ**m particles or cells, and *C_total_* denotes all 10 **μ**m particles or cells delivered to droplets.

## Results and Discussion

A first requirement for the high throughput analysis of single cells in droplets is the sustained and homogeneous delivery of high concentration bead and cell suspensions. This is especially the case for the large and dense reporter beads which rapidly sediment yet are also fragile. Initially this challenge was solved using perpetual sedimentation (rotation of a horizontal syringe to produce bead orbits, effectively making them neutrally buoyant)^23^. Then, to aid broader uptake in conventional cell biology labs we instead used a vertically orientated syringe with a powerful electromagnet to gently rotate a disc magnet: The disc magnet disperses beads and cells in all directions, sideways for mixing, upwards to return by gravity and downwards to exit. Both approaches enabled the sustained delivery of cell and particle suspensions suitable for investigating Dean entrainment effects (**SI Fig. 3**).

To gain a first understanding of entrainment we employed monodisperse 30 **μ**m polystyrene particles to represent ToyoPearl reporter beads and monodisperse 10 **μ**m polystyrene particles to represent mammalian cells. The emergence of particle entrainment, from disordered to periodic spacing was observed with a 600k/mL 30 **μ**m particle suspension using a 100 mm/s mean flow velocity (*Re_p_* 0.9, **SI Fig. 4A**). Entrainment requires concentrated suspensions with particle train length increasing and inter-particle pitch decreasing with concentration. At 1 million/mL (1.4% volumetric fraction (*vf*)) a median inter-particle pitch of 75 **μ**m was produced (**Figure 2A**). This equates to 2.5*D* (*D = diameter/pitch*) arrangements associated with higher *Re_p_* numbers^15^ and is attributed to the high volume fraction suspension and prolonged inertial transport (100 mm) supplemented with secondary Dean flows. The concentration could be extended to 1.5 million/mL (2.1% *vf*), but above this crowding effects result in loss pf periodicity (**SI Fig. 4B**). Entrainment of the 10 **μ**m polystyrene (*Ū* = 100 mm/s, *Re_p_* 0.2) particles was also concentration-dependent, with striking ordering observed at 6 million/mL (0.3% *vf*, **Figure 2B**) producing a median *5D* pitch. At these moderate particle concentrations, the increased gaps between trains extends the pitch, and pitch variability. Higher volumetric fractions are feasible. However, in the context of cell processing, such concentrations are unsuitable for maintaining cell viability and promote cell clustering.

**Figure 2.**
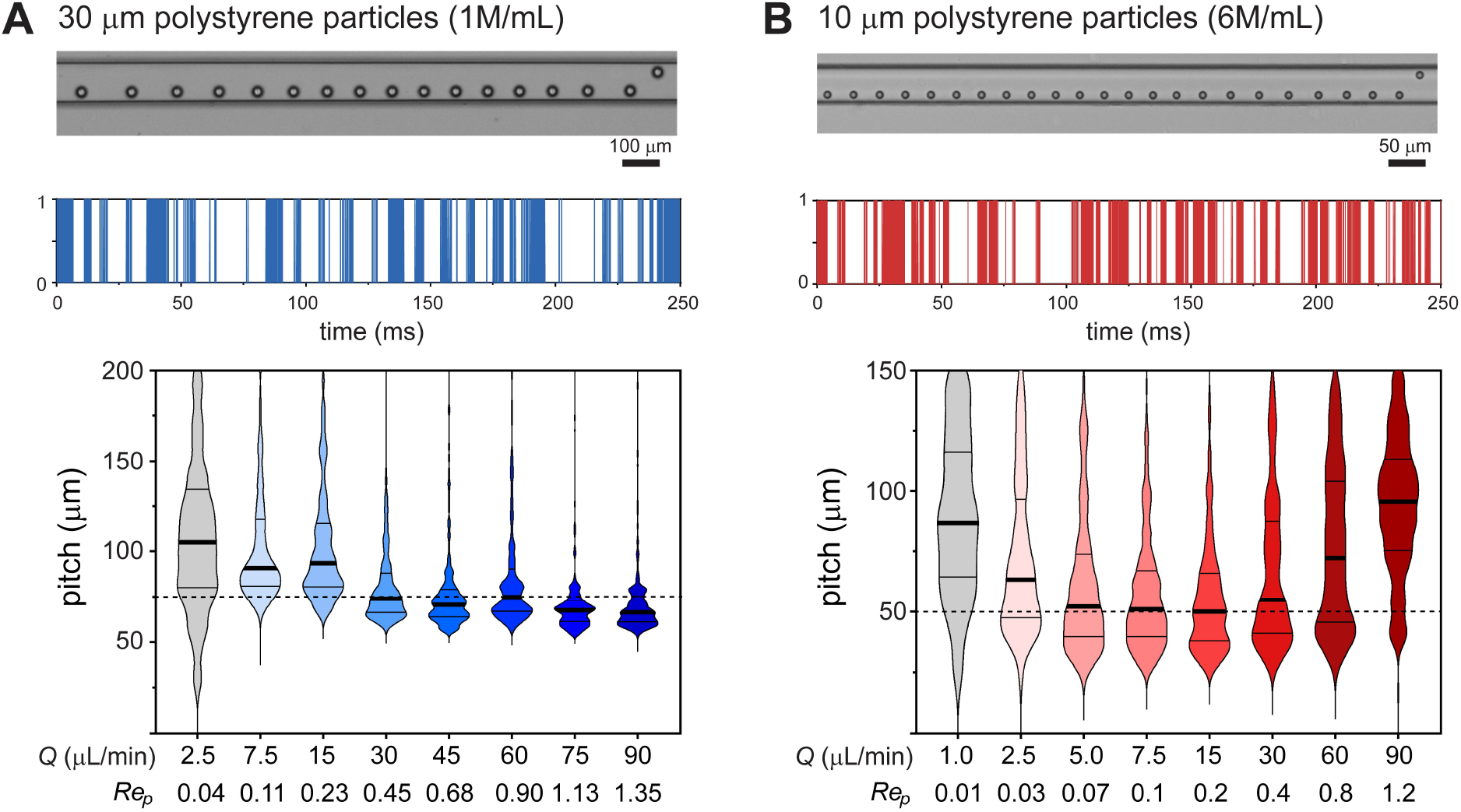
Scalable particle entrainment. Exemplary frames of 30 μm particle **(A)** and 10 μm particle **(B)** entrainment with a mean flow velocity of 100 mm/s. Entrainment intensity profile with distance translated to time of a 250 ms imaging segment illustrating gap and train length variability. Violin plots of the velocity dependence (flow rate and *Re_p_*) of the particle pitch distribution represented as median, 25^th^ and 75^th^ percentiles and data extremities with cut-offs at 200 μm for 30 μm particles and 150 μm for 10 μm particles (*n* >2,000 particles per velocity condition). The 2.5*D* and 5*D* inter-particle pitch predictions are indicated with dashed lines. Grey violin plots denote random particle distributions, without entrainment.

The velocity dependence was investigated using flow rates ranging from 1–90 **μ**L/ min, producing a velocity range of 1.7–150 mm/s (*Re_p_* 0.015–1.35, *De_max_* 0.04–3.75) for the 30 **μ**m particles and 6.7–600 mm/s (*Re_p_* 0.01–1.2, *De_max_* 0.06–5.30) for the 10 **μ**m particles. 30 **μ**m particles were tightly entrained with a 2.5*D* pitch using a 30–90 **μ**L/min flow range (**Figure 2A**), and 10 **μ**m particles entrained with a 5*D* pitch using a 5–30 **μ**L/min flow range (**Figure 2B**). At lower flow rates entrainment quality diminishes, ultimately leading to randomly distributed particles with different velocities.

The inter-particle pitch results can be used to predict 30 and 10 **μ**m particle volume limits for effective single bead and single cell co-encapsulation: Using the 25^th^ percentile data, volumes below 650 pL are needed for 30 **μ**m particles and volumes below 100 pL for 10 **μ**m particles to obtain efficient co-encapsulations. This requires different bead and cell flow rates. Using these single particle volume packets as a guide we introduced flow conditions for the generation of 600 pL droplets. To identify optimal 30 **μ**m particle (referred to as ‘beads’) and 10 **μ**m particle (referred to as ‘cells’) input volumes for each droplet, a volume ratio scaling experiment was undertaken: The ‘bead’ and ‘cell’ flow rates (*Q_bead_* and *Q_cell_*) were differentially modulated to produce volume ratios ranging from 1 to 15 (300_*bead*_+300_*cell*_ pL to 562_*bead*_+38_*cell*_ pL) while satisfying the requirements for effective Dean entrainment. The co-encapsulation results are compared with theoretical results from the joint probability distribution (JPD) in **Figure 3A**. With large ‘cell’ volumes dual Dean entrainment has a substantially reduced single ‘cell’ and ‘bead’ coupling frequency due to entrainment increasing the ‘cell’ multiplet rate. At a volume ratio of 10 (545+55 pL) a transition occurs in which entrainment becomes beneficial, with the noise (‘cell’ multiplets) dropping below the theoretical JPD value. As the volume ratio further increases, the ‘cell’ multiplet rate tends to zero, while the signal (single ‘cell’ capture rate) remains similar to the JPD results, allowing high throughput and extreme signal-to-noise processing.

**Figure 3.**
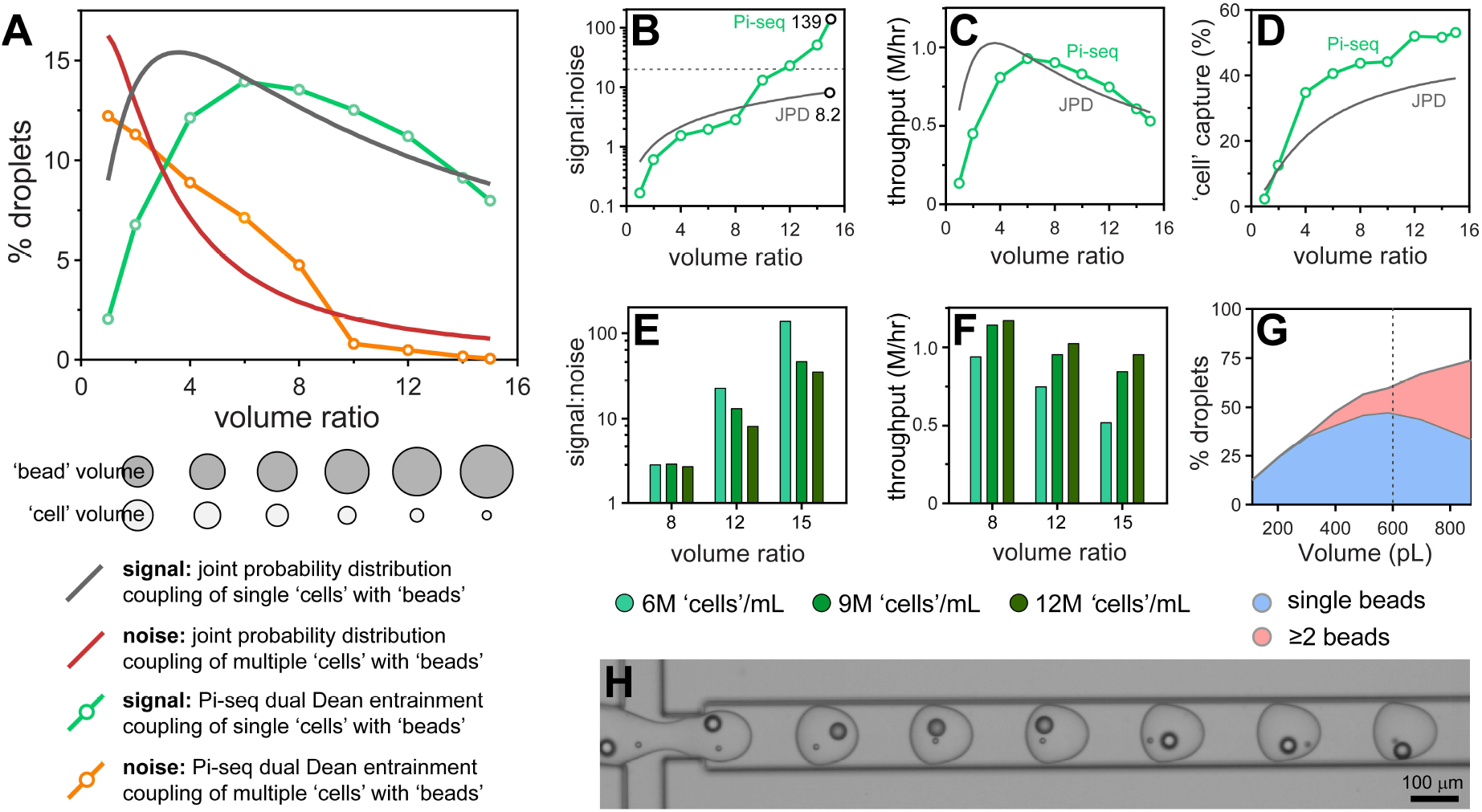
Volume ratio effects on polystyrene particle co-encapsulation. Droplet volumes were standardized at 600 pL and the ratio between cell and bead volumes modulated between 1:1 and 1:15 (300:300 pL to 38:562 pL). The monodisperse 10-μm particles are used to approximate cells and denoted as ‘cells’, and monodisperse 30-μm particles approximating the ToyoPearl beads denoted as ‘beads’. Pirouette-seq (Pi-seq) co-encapsulation results are plotted with the theoretical JPD for given volume ratios with a ‘bead’ concentration of 1.2 million/mL and a ‘cell’ concentration of 6 million/mL. The percentage of droplets producing a signal (BC, BBC) and noise (BCC, BBCC) was obtained by analyzing ~3,700 droplets per condition **(A)**. The volume ratio effect on signal-to-noise with the *S:N* 20 threshold indicated using a grey dashed line **(B)**, throughput **(C)** and capture rate **(D)**. Increasing the ‘cell’ concentration from 6 million/mL to 9 million/mL and 12 million/mL reduces the signal-to-noise **(E)** while increasing the throughput **(F)**. Data obtained by analyzing ~1,850 droplets per condition. Increasing the droplet volume from 110 to 890 pL results in higher proportions of droplets containing multiple ‘beads’ **(G)**. Frame documenting an ideal single ‘cell’ and single ‘bead’ co-encapsulation sequence using a volume ratio of 1:12 **(H)**.

To appreciate the different performance metrics the signal-to-noise, throughput and capture efficiency are compared with JPD predictions in **Figures 3B-D**. These results demonstrate the merits of dual Dean entrainment; with a volume ratio of 15 the signal-to-noise is 139 (0.7% multiplets), 17-fold higher than random delivery, along with a throughput of 0.5 million/hr and a capture efficiency >50%. The gains in capture efficiency above the JPD prediction result from bead entrainment that produces higher numbers of droplets containing a ‘bead’. The 6 million/mL ‘cell’ concentration has a low volumetric fraction (0.3%), indicating scope for higher cell concentrations. However, higher concentrations introduce localized crowding effects such that gains in throughput are at the expense of the signal-to-noise (**Figures 3E,F**). To increase the capture efficiency larger droplet volumes were considered. This allows >70% of droplets to contain a bead, enabling >70% of ‘cells’ to be captured (see **Figures 3G**). However, this would require lower ‘cell’ flow rates, insufficient for effective entrainment. The **SI video** documents dual Dean entrainment for the co-encapsulation of periodically spaced ‘cell’ and ‘bead’ trains into droplets. Ideal results are shown in **Figure 3H** and typical results in **SI Fig. 5**.

Given the promising performance with polystyrene particles we next sought to answer whether dual Dean entrainment can be effectively applied to the co-encapsulation of ToyoPearl beads (unfunctionalised ChemGene beads used in the Drop-seq protocol) with mammalian cells. Both ToyoPearl beads (ø34.1±3.3**μ**m) and human HEK293 cells (ø14.3±1.5 **μ**m) were effectively entrained at the same concentrations used for the polystyrene particle experiments (**SI Fig. 4C and 6**). In addition, the shear flow conditions rapidly disperse cell clusters into trains of single cells, potentially representing a means for sample disaggregation (**SI Fig. 7**). The bead and cell flow rate dependent pitch distributions closely followed those obtained for the 10 and 30 **μ**m polystyrene particles, although the HEK293 cells produced a 3.5*D* median pitch (10 **μ**m particles = 5*D*). The results from a volume ratio scaling co-encapsulation experiment are documented in **Figure 4A-E**. Noise reduction beneath the JPD prediction occurred later with a flow ratio of 16 and stabilized with a flow ratio ≥20, overall resulting in a modest signal-to-noise (24; 4% multiplet rate). Throughput was maintained at ~0.5 million cells/hour, and the capture efficiency was extended to ~70% by using a 1.5 million/mL ToyoPearl bead concentration. Performance was corroborated by repeating the experiment with a THP cell line (**SI Fig. 8**). These smaller cells (ø11.0±2.7 **μ**m) eliminate size effect interpretations.

**Figure 4.**
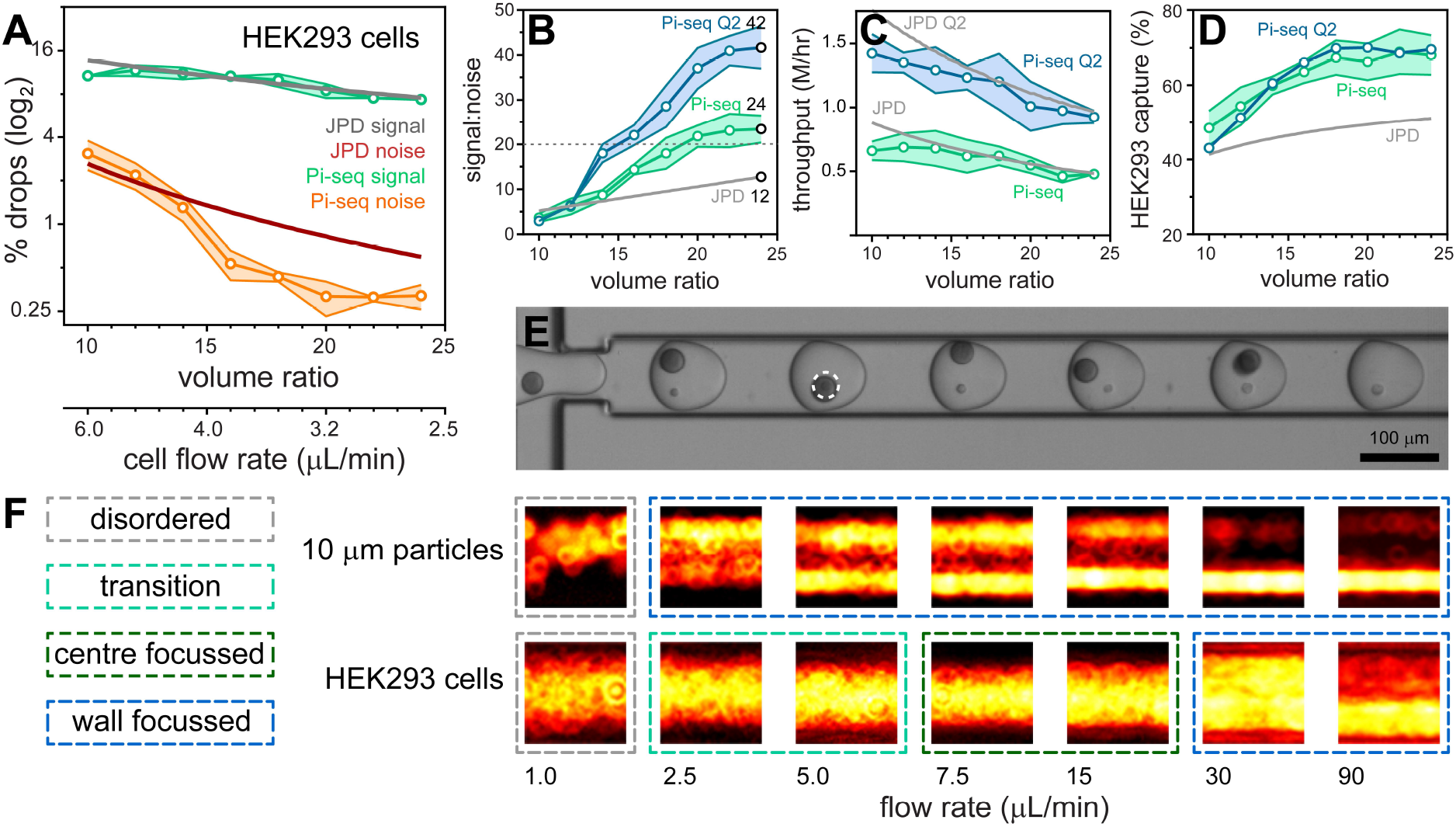
Volume ratio effects on the co-encapsulation of ToyoPearl beads with HEK293 cells. Droplet volumes were standardized at 600 pL and the ratio between cell and bead volumes modulated between 1:10 and 1:24 (55:545 pL to 24:576 pL). Pirouette-seq (Pi-seq) co-encapsulation results are plotted with the theoretical JPD for given volume ratios with a ToyoPearl bead concentration of 1.5 million/mL and a HEK293 cell concentration of 6 million/mL. The percentage of droplets producing a signal (BC, BBC) and noise (BCC, BBCC) was obtained by analyzing >3,500 droplets per condition (*n*=3 experiments) **(A)**. The volume ratio effects on signal-to-noise with the *S:N* 20 threshold indicated using a grey dashed line **(B)**, throughput **(C)** and capture rate **(D)** are plotted for standard (Pi-seq; green) and double (Pi-seq Q2; blue) aqueous flow rates. Frame documenting an efficient single HEK293 cell and single bead co-encapsulation sequence using a volume ratio of 1:16 **(E)**. The 10 μm polystyrene particles and HEK293 cells have different flow rate-dependent inertial focusing behaviors **(F)**.

We sought to understand the late onset of noise reduction and absence of exponential signal-to-noise scaling with increasing volume ratio. The 10 **μ**m polystyrene particles and HEK293 cells produce equivalent pitch minima (~20 **μ**m), but distinctly different focusing behavior (**Figures 2B** and **4F, SI Figs. 6** and **9**): The solid 10 **μ**m particles are wall-focussed for all flow rates excepting 1 **μ**L/min in which focusing collapses, producing random cross-channel positions. In stark contrast, cells are deformable with inner wall focusing only occurring above 30 **μ**L/min (*Re_p_*>1, **SI Fig 7D**), below which HEK293 cells become entrained within streamlines towards the channel centre^11, 24^ and transition towards random, unfocussed positions at the flow rates required to produce high volume ratios. Cell transport within different streamlines, with different velocities, increases the probability of cells arriving together at the droplet generation junction. To achieve higher signal-to-noise sample processing, higher *Re_p_* flow conditions are needed for effective cell entrainment. We investigated the upper limits of the dripping droplet formation regime: Aqueous flow rates can be doubled in combination with a 240 **μ**L/min oil flow rate while retaining a 600 pL droplet volume. Repeating the volume ratio scaling experiment using these elevated flow conditions improved the signal-to-noise to 42 (2.3% multiplets) with the benefit of doubling the throughput to ~1 million cells/hour (~15,000/minute) while retaining the ~70% capture efficiency (**Fig. 4B-D**). Higher flow regimes enter a jetting regime with bead-triggering producing higher droplet generation rates for even higher throughput processing.^25^ However, bead-triggered droplet formation in the jetting regime increases droplet polydispersity, at odds with precisely defined volumes required to effectively co-encapsulate single cells by entrainment.

Pirouette-seq out-performed commercial and entrainment-based single cell and reporter bead coupling methods (**SI Fig. 10, SI Table 2**). Alternative approaches bypass the Poisson-dictated multiplet rate problem by pre-indexing cells by labelling membranes (MULTI-seq^26^) or transcriptomes (sci-seq^27, 28^, SPLiT-seq^29^ and scifi-RNA-seq^30^). The scifi-seq method was used to allow droplet ‘super-loading’, demonstrating a throughput of >150,000 nuclei per 10X channel (>500,000 nuclei/hour). Each technique has its own deficiencies, such as lengthy procedures, labels being exchanged, cell losses during labelling and volume limitations restricting analyses to nuclei (foregoing the information content from the rest of the cell). Nevertheless, substantial improvements can readily be anticipated in these and other approaches for single cell indexing. Indeed, the current Pirouette-seq prototype represents a blueprint for future iterations incorporating refinements to microfluidic dimensions allowing, for instance, improved focusing (**SI Fig. 4D**) and operation at higher *Re_p_* numbers for enhanced signal-to-noise processing. In general, these technologies forecast the routine undertaking of large-scale experiments that will become feasible as dramatic cost savings begin to emerge from innovations in sequencing^31^.

The coupling efficiencies enabled by Pirouette-seq allow other analytical scenarios to be envisaged, such as experiments requiring cells to be rapidly processed to prevent transcriptome remodeling, those involving different beads reporting different biological dimensions, or a bead to perturb the cell and another bead to report biological outcomes. For example, Pirouette-seq offers the potential for screening genetically-encoded bead-based compound libraries without exhaustive passes to ensure library coverage. Here, the ability of Dean entrainment to process high concentrations of solid beads allows the repertoire of solid phase synthesis methods to be used in library construction. Overall, the coupling efficiencies lend Pirouette-seq to large-scale experiments that were previously impractical.

## Conclusions

Pirouette-seq combines cell and bead entrainment to bypass the limitations of the joint probability distribution during droplet co-encapsulations. This produces profound gains in performance, achieving extreme throughput combined with an enhanced signal-to-noise while capturing the majority of cells. The approach has broad-reaching potential, enabling cellular systems to be comprehensively profiled in health, disease and in response to perturbation.

## Supporting information

CoEncapsulation Video

## Acknowledgements

We are indebted to Deborah Mackay for her support. We also thank Gessa Sugiyarto for preparing cells. The project was made possible by financial contributions from the Medical Research Council (JB and MSJ: MC_PC_15078), a CRUK & Royal College of Surgeons of England Advanced Clinician Scientist Fellowship (JH and TU), an EPSRC CASE studentship (LBE), GSK (AFV, ARCP006668), a Sir Henry Dale Wellcome Trust Fellowship (MEP; 109377/Z/15/Z) and a John Goldman Fellowship (MRZ; 2016/JGF/0003).

## Author Contributions

JH, LBE, JB, AFV, SIRL, BS, MSAJ, RP, PSS, RCGS, MRZ and JW undertook the experiments, BDM, MRZ, MEP, TU and JW supervised the project, and JW wrote the manuscript, with co-authors reviewing the manuscript.

