## Supplementary Information for "Dual Dean entrainment with volume ratio modulation for efficient co-encapsulation: Extreme single-cell indexing"

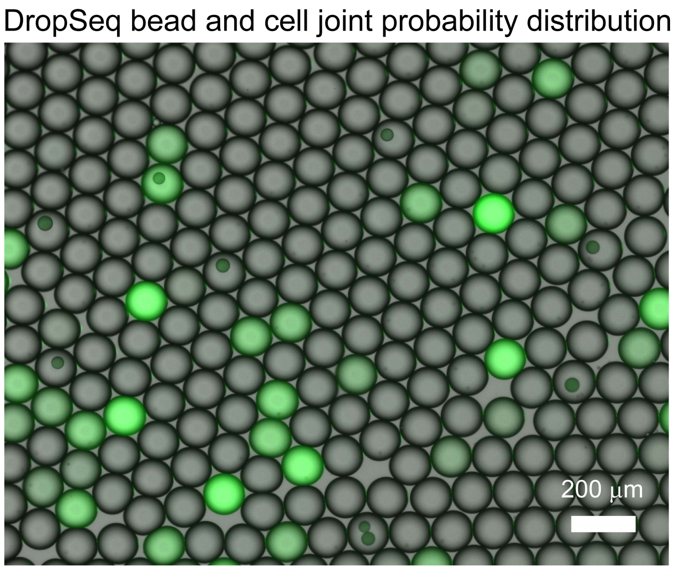


**SI Figure 1.** Illustration of the co-encapsulation improbability using standard Drop-seq microfluidic conditions[1] to pair ChemGene beads with calcein AM loaded cells that are lysed upon encapsulation.

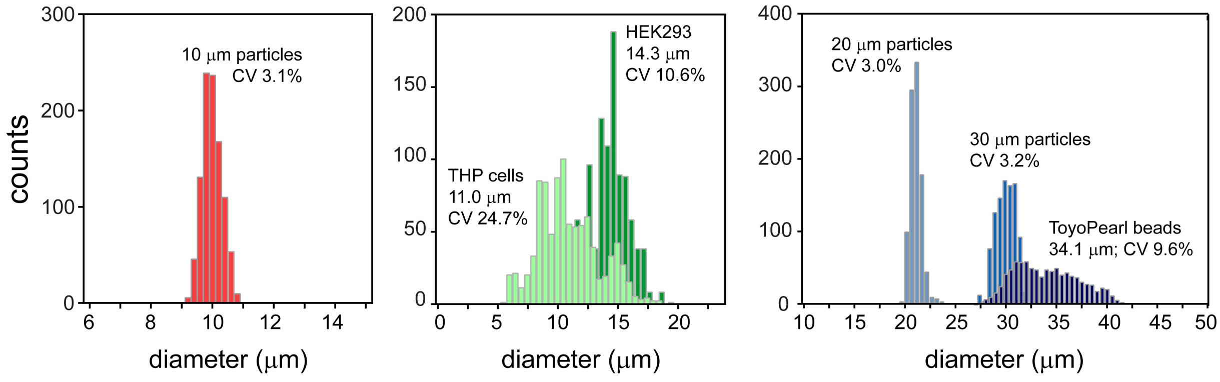


**SI Figure 2:** Polystyrene particle, mammalian cell and ToyoPearl bead diameter distributions (*n*=1,000).

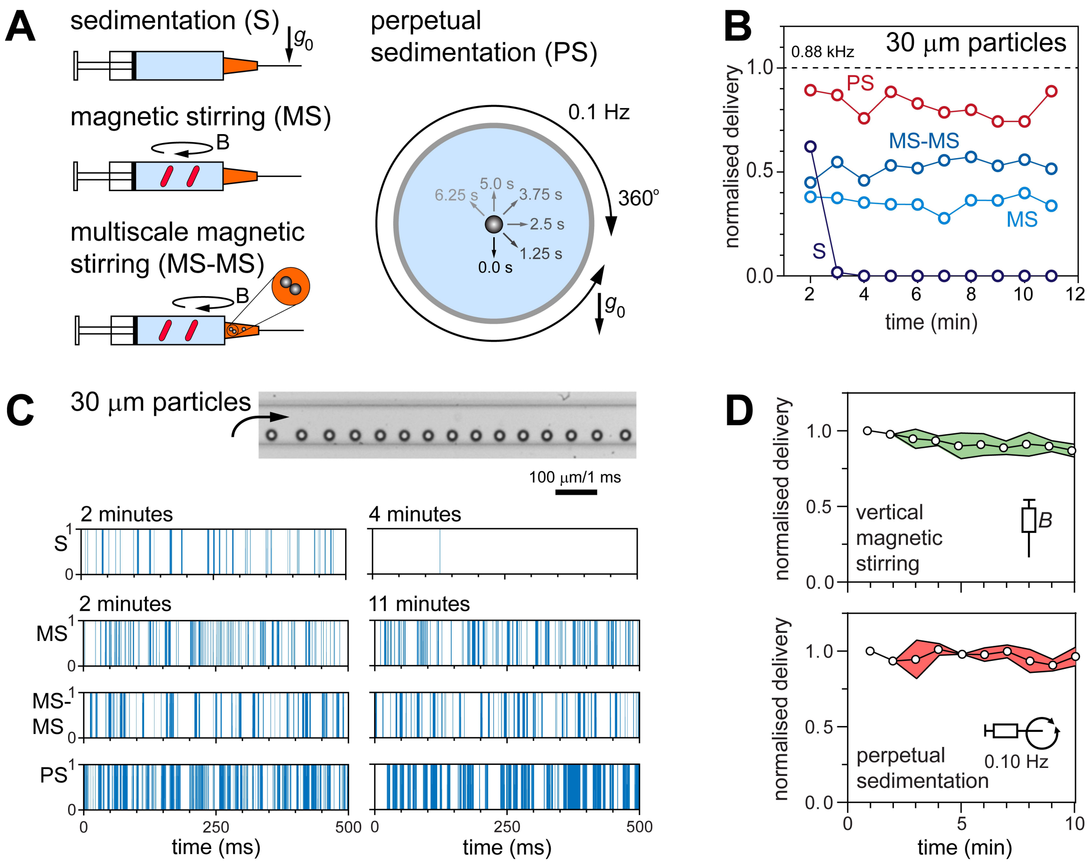


SI Figure 3. *Continuous particle delivery.* Particle delivery arrangement used to measure sedimentation effects (S, control) were compared with magnetic stirring (MS) using two 7 mm magnetic bars, multi-scale magnetic stirring (MS-MS) using an additional three 1-mm-diameter stainless steel (AISI 420) ball bearings in the Luer-lock tip, and cross-section of the syringe barrel with 360° rotation and reversal to effect perpetual sedimentation (PS) **(A)**. Delivery frequency 11-minute time course for 30-μm-diameter particles. Data is normalised as a function of the 0.88 kHz theoretical frequency **(B)**. Frame of entrained particles and delivery frequency traces (with distance translated to time) recorded over 500 ms at the beginning and end of each time course **(C)**. Both perpetual sedimentation and vertical syringe orientation with a disc magnet actuated using a low rpm (25–30 a.u.) drive magnet maintain particles in suspension for sustained delivery to the spiral Dean entrainment microchannel **(D)**. Reproduced from the original RSC *Lab on a Chip* article[2].

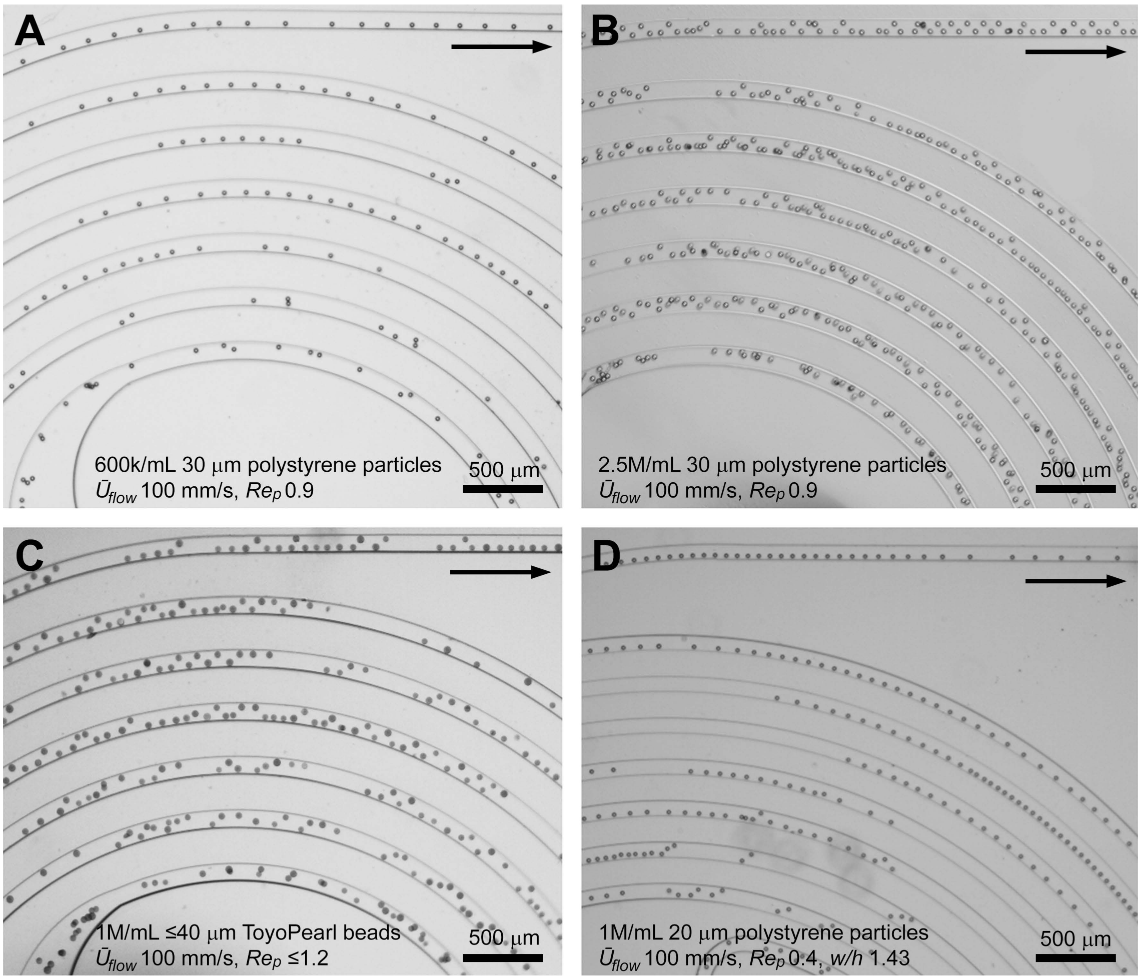


SI Figure 4: *Entrainment characteristics.* Emergence of 30 μm polystyrene particle trains from a disordered state at the spiral entrance to periodic spacing at moderate 600k/mL concentrations (A). An overcrowded particle concentration of 2.5 million/mL results in a near-continuous particle train, with particle alternation at either side of the channel, loss of periodicity and particle contacting (B). Polydisperse (CV 9.6%) ≤40 μm ToyoPearl particle trains experience alternation at 1 million/mL concentrations (C). Changing the channel aspect ratio to 1.43 (*w* 100 μm; *h* 70 μm) and using smaller, 20 μm particles removes particle alternation patterns at 1 million/mL concentrations (D).

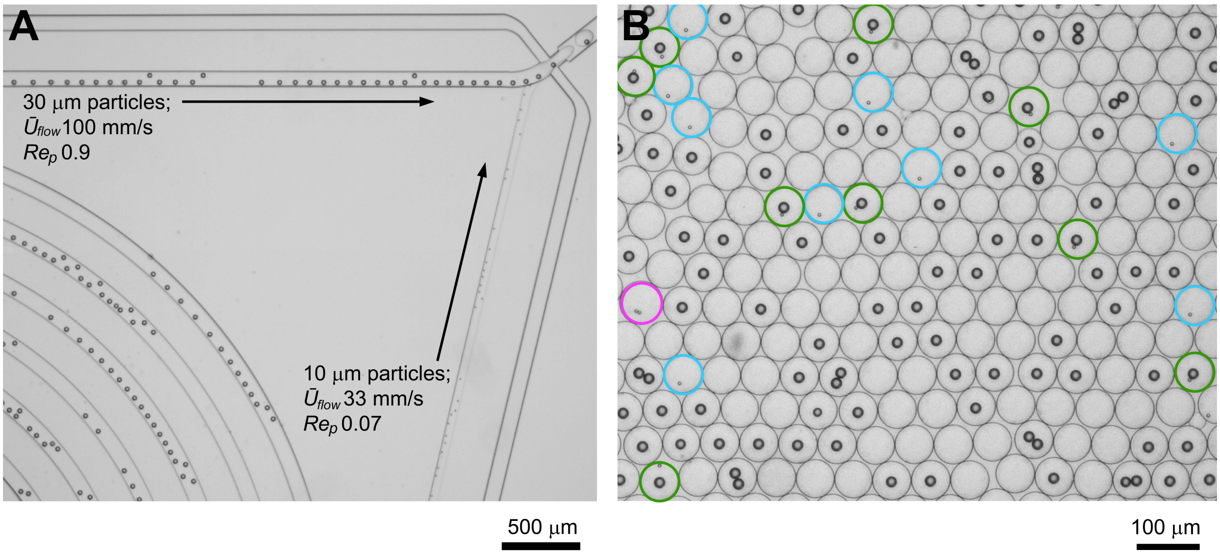


**SI Figure 5.** *Co-encapsulation.* Convergence of entrained 10 μm and 30 μm particles upstream of droplet encapsulation **(A)** involving flow rates of 5 μL/min for 10 μm particles and 60 μL/min for 30 μm particles producing a volume ratio of 1:12. Image illustrating encapsulation results, with green rings identifying single 'cell' and single 'bead' co-encapsulation events, blue rings identifying single 'cell' encapsulation events and the purple ring identifying a 'cell' doublet encapsulation event **(B)**.

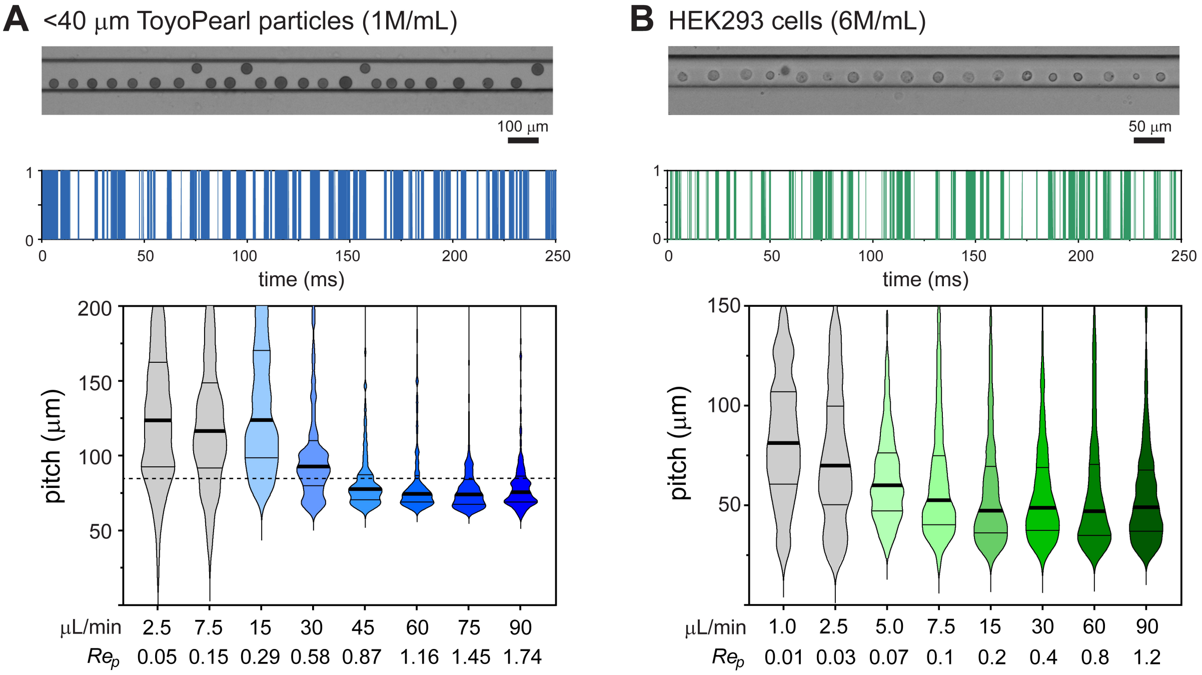


**SI Figure 6.** ToyoPearl bead **(A)** and HEK293 cell **(B)** entrainment. Exemplary frames of ≤40 μm ToyoPearl particle and HEK293 cell (Ø14.3±1.5 μm) entrainment with a mean flow velocity of 100 mm/s. Entrainment intensity profile from a 250 ms imaging segment illustrating gap and train length variability (distance translated to time). Violin plots of the velocity dependence (flow rate and $Re_p$) of the bead and cell pitch distribution represented as median, 25th and 75th percentiles and data extremities with cut-offs at 200 μm for ToyoPearl beads and 150 μm for HEK293 cells (*n* >2,500 beads or cells per velocity condition). The 2.5$D$ inter-particle pitch prediction for the ToyoPearl beads is shown with a dashed line. Grey violin plots denote random particle distributions, without entrainment.

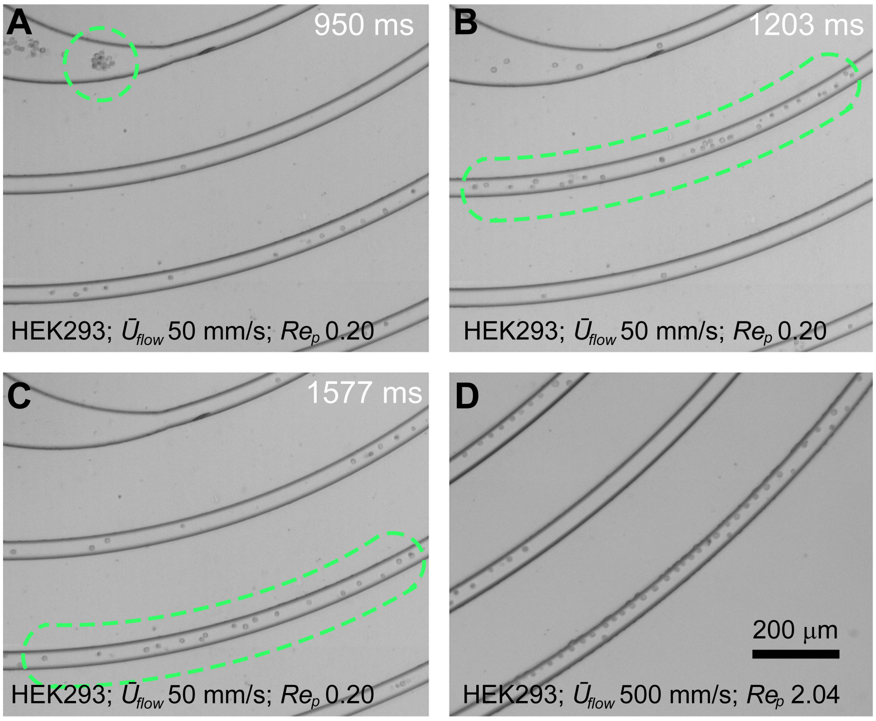


**SI Figure 7.** HEK293 cell clusters are rapidly disaggregated and dispersed into cell trains at a flow rate of 7.5 μL/min (50 mm/s, $Re_p$ 0.2, **A-C**). HEK293 cells predominantly undergo inner wall focusing at high flow rates (7.5 μL/min, 500 mm/s, $Re_p$ 2.04, **D**).

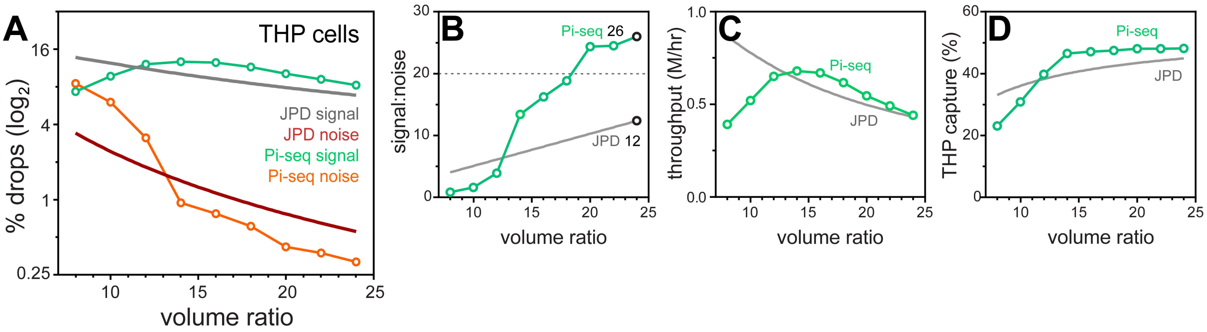


**SI Figure 8.** Volume ratio effects on the co-encapsulation of ToyoPearl beads with THP cells. Droplet volumes were standardized at 600 pL and the ratio between cell and bead volumes modulated between 1:8 and 1:24 (67:533 pL to 24:576 pL). Pirouette-seq (Pi-seq) co-encapsulation results are plotted with the theoretical JPD for given volume ratios with a 1.2 million/mL ToyoPearl bead concentration and a 6 million/mL THP cell concentration. The percentage of droplets producing a signal (BC, BBC) and noise (BCC, BBCC) was obtained by analyzing >3,500 droplets per condition **(A)**. The volume ratio effect on signal-to-noise with the *S:N* 20 threshold indicated using a grey dashed line **(B)**, throughput **(C)** and capture rate **(D)**.

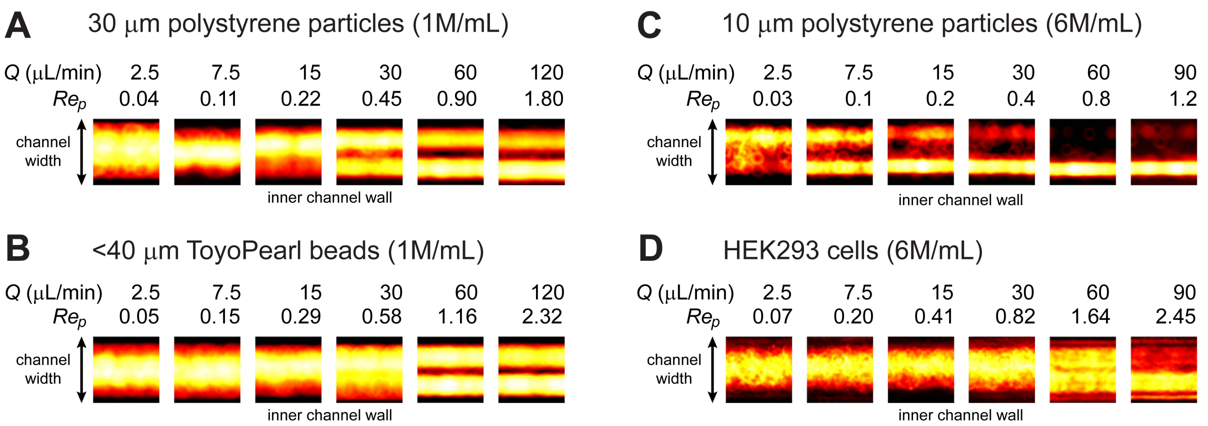


**SI Figure 9.** The solid 30 μm polystyrene particles **(A)** and ≤40 μm ToyoPearl beads **(B)** have the same velocity-dependent focusing behavior. The solid 10 μm polystyrene particles maintain wall focusing throughout the velocity range, with random, non-entrained, particle distributions only emerging at low flow velocities, $Re_p$ 0.033 **(C)**. The larger HEK293 cells (Ø14.3±1.5 μm) have different velocity-dependent focusing characteristics; wall focused positions at $Re_p$ ≥1.0, moving to channel centre focusing positions at $Re_p$ 0.5 and then transitioning to unfocussed random positions at $Re_p$ <0.02 **(D)**. The deformable nature of cells transported within inertial flows explains this difference[3-5].

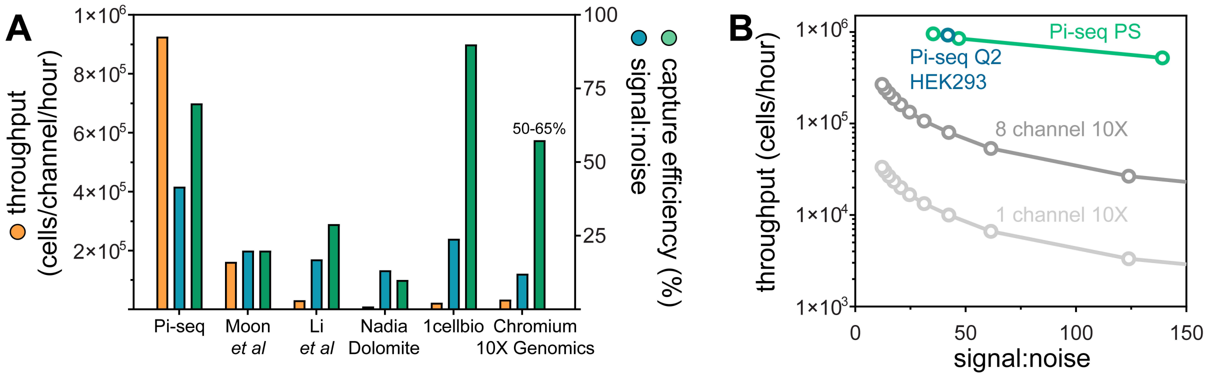


**SI Figure 10.** Pirouette-seq (Pi-seq) performance compared with commercial co-encapsulation systems and academic systems involving inertial ordering **(A)**. Pi-seq compared with single channel and 8 channel Chromium 10X Genomics processing **(B)**. Pi-seq data from coupling 10 and 30 μm polystyrene particles (green) and with flow rate doubling for coupling HEK293 cells with ToyoPearl beads (blue).

#### Table 1. Flow conditions for volume ratio scaling

Flow conditions for modulating cell and bead droplet volumes. $\bar{U}_{flow}$ denotes mean flow velocity, and $Re_p$ the particle Reynolds number obtained from $Re_p=Re(a/D_h)^2$, where $Re=\rho D_h \bar{U}/\eta$, $\rho$ is the fluid density, $\eta$ dynamic viscosity and $R$ is the spiral channel minimum radius (1.6 mm). The $Re_p$ values are calculated using a 10 μm cell diameter and a 30 μm bead diameter. The fluoro-oil flow rate was kept constant at 165 μL/min. Higher $Re_p$ operation was achieved by doubling the aqueous inputs in combination with a constant 240 μL/min fluoro-oil flow rate.

| Volume Ratio | Cell Flow Rate | Cell $\bar{U}_{flow}$; $Re_p$ 10 μm | Cell Droplet Contribution | Bead Flow Rate | Bead $\bar{U}_{flow}$; $Re_p$ 30 μm | Bead Droplet Contribution |
|---|---|---|---|---|---|---|
| 1:1 | 33 μL/min | 220 mm/s; 0.44 | 300 pL | 33 μL/min | 55 mm/s; 0.50 | 300 pL |
| 1:2 | 22 μL/min | 147 mm/s; 0.29 | 200 pL | 44 μL/min | 73 mm/s; 0.66 | 400 pL |
| 1:4 | 13 μL/min | 87 mm/s; 0.17 | 120 pL | 52 μL/min | 87 mm/s; 0.78 | 480 pL |
| 1:6 | 9.5 μL/min | 63 mm/s; 0.13 | 86 pL | 57 μL/min | 95 mm/s; 0.86 | 514 pL |
| 1:8 | 7.5 μL/min | 50 mm/s; 0.10 | 67 pL | 60 μL/min | 100 mm/s; 0.90 | 533 pL |
| 1:10 | 6 μL/min | 40 mm/s; 0.08 | 55 pL | 60 μL/min | 100 mm/s; 0.90 | 545 pL |
| 1:12 | 5 μL/min | 33 mm/s; 0.067 | 46 pL | 60 μL/min | 100 mm/s; 0.90 | 554 pL |
| 1:14 | 4.5 μL/min | 30 mm/s; 0.060 | 40 pL | 63 μL/min | 105 mm/s; 0.95 | 560 pL |
| 1:15 | 4 μL/min | 27 mm/s; 0.053 | 38 pL | 60 μL/min | 100 mm/s; 0.90 | 562 pL |
| 1:16 | 4 μL/min | 27 mm/s; 0.053 | 35 pL | 64 μL/min | 107 mm/s; 0.96 | 565 pL |
| 1:18 | 3.5 μL/min | 23 mm/s; 0.047 | 32 pL | 63 μL/min | 105 mm/s; 0.95 | 568 pL |
| 1:20 | 3.2 μL/min | 21 mm/s; 0.043 | 29 pL | 64 μL/min | 107 mm/s; 0.96 | 571 pL |
| 1:22 | 2.9 μL/min | 19 mm/s; 0.039 | 26 pL | 64 μL/min | 107 mm/s; 0.96 | 574 pL |
| 1:24 | 2.7 μL/min | 18 mm/s; 0.036 | 24 pL | 64.8 μL/min | 108 mm/s; 0.97 | 576 pL |

#### Table 2. Single-cell RNA-Seq co-encapsulation method comparison

Key performance metrics comparison commercial platforms based on droplet microfluidics and nanowell array approaches and academic systems involving entrainment. Co-encapsulation metrics from the described dual Dean entrainment system in blue, co-encapsulation metrics considering random delivery (*i.e.* without entrainment) for input conditions based on the joint probability distribution in green and explanation of the normalisation of throughput per channel per hour in grey text. Data obtained March 2021.

| Class | Method | Throughput per Channel | Multiplet Rate | Capture Efficiency |
|---|---|---|---|---|
| *Droplet microfluidics with entrainment* | Pirouette-seq (1:24) | 927,000/hr | 2.3% | 70% |
| | JPD predictions | 968,000/hr | 13% | 51% |
| | Bead Inertial Ordering[6] | 162,000/hr | ~4% | 20% |
| | JPD predictions | 168,000/hr | 3% | 21% |
| | Bead+Cell Entrainment[7] | 31,100/hr | >5% | 29% |
| | JPD predictions | 24,500/hr | 5% | 23% |
| *Droplet microfluidics with solid beads* | DropSeq[1] | 10,000/hr | ~5% | ~10% |
| | Nadia (Dolomite)[8] | 18,750/hr<br>6250/channel in 20 mins;<br>8 channels/cartridge | 6–8% | ~10% |
| | ddSEQ (BioRad)[8, 9] | 4,500/hr<br>$375_{max}$/channel in 5 mins;<br>4 channels in parallel | ~6% | 3% |
| *Droplet microfluidics with hydrogel beads* | InDrop[8, 10, 11] | 4,000–12,000/hr | ~4% | >90% |
| | 1CellBio[8] | 22,500/hr<br>5,000/channel;<br>8 channels/cartridge | ~4% | >90% |
| | 10X Genomics[12] | 33,300/hr<br>10,000/channel in 18 mins;<br>8 channels/cartridge | 7.6% | 50–65% |
| *Nanowell microarrays with solid beads* | Cyto-Seq[13] | ~1500/hour<br>2,300 indexed per array,<br>1.5 hour processing | 5.2%<br>($\lambda$<0.1) | 14.1% |
| | Seq-Well[14] | 5,300/hour<br>20,000 cells loaded<br>with a 3 hour protocol | 4.8% | 80% |
| | Microwell-Seq[15] | 40,000/hour<br>10,000/array in 15 mins | 1.2% | 10% |

#### Appendix I: Instructions for establishing particle entrainment

**Context:** Entrainment is achieved by transporting high-concentration particulate suspensions within channels of similar-scale dimensions. Such conditions are prone to microchannel clogging resulting from the production of volume packets saturated with particles (*e.g.* >5% v/v). A number of factors can generate these particle-saturated volumes; (a) the use of large and/or dense particles that are prone to sediment, (b) the accumulation of particles at air-liquid interfaces, and (c) the arrival of environmental fibres or particles within the microchannel that are sufficiently large to become lodged. These issues can be readily addressed to enable the reliable and sustained processing of high-concentration suspensions in moderate and high velocity microfluidic formats:

| | |
|---|---|
| ***Filtration*** | Filter all solutions prior to the addition of particles and cells using, for example, using a 0.22 μm filter (Millex®, Millipore). |
| ***Syringe Orientation*** | Particles should be delivered from a vertically-orientated syringe, pointing downwards and elevated above the microfluidic device using for example a jack to raise the syringe pumps above the microscope stage. The tubing connection should be appropriately short to eliminate horizontal sections where particles can sediment. Alternatively, perpetual sedimentation configurations can be used[1]. |
| ***Magnetic Stirring*** | Rapidly settling particles require stirring within the syringe. We recommend using a disc magnet actuated by a powerful driving electromagnet such as the Multi Stirrus™ (V&P Scientific) system. Such powerful systems allow relatively slow disc magnet rotation, ensuring particles are retained in suspension without being damaged. This is especially important for the brittle ToyoPearl/ChemGene beads. |
| ***Remove Air Pockets*** | Air is removed from the channels by flooding the microfluidic device with fluoro-oil. Critically, air should also be removed from the inlet ports by closing the droplet exit port using a 1-mm-diameter stainless steel pin or similar. Fluoro-oil is diverted to the 100x100 μm Dean spiral channel for beads, filling the inlet port to the brim. Then, stop the fluoro-oil flow and remove the steel pin in readiness for inserting tubing. |
| ***Coupling Tubing*** | First, ensure air bubbles are removed from the tubing. To prevent particles accumulating at the air–liquid interface at the tubing terminus particle suspension flows are established at substantial (*e.g.* 50 μL/min) flow rates before inserting tubing within the microfluidic ports. Both bead and cell flows are run for 10's of seconds into microcentrifuge tubes to ensure flows are fully established (and reagents can be recycled) and then, during continuous flow, tubing is inserted into the oil-filled inlet ports. Once inserted the fluoro-oil flow is immediately started to assist beads exiting the device within droplets. Cell and bead entrainment is established in <1 minute after which cell and bead inputs are set to the desired flow rates. |

### Appendix II: Particle Pitch MATLAB Code

#### Code Overview

This code measures the pitch and delivery frequency of entrained particles or cells flowing in a linear microfluidic channel. Using ImageJ the original video file is first transformed to a stacked TIFF, with the use of rotation and cropping tools to define the image analysis window.

To increase the signal to noise ratio of a particle/cell compared to the channel background, a background image of the channel with no particles present is subtracted from each image in the sequence (**Figure 1**).

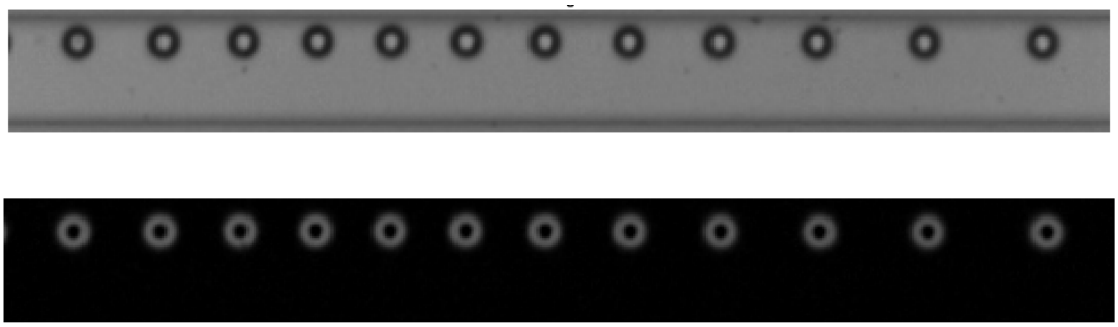

*Figure 1: Raw image of particles flowing in a microchannel (top). Background subtracted image (bottom).*

The pixel intensity for each image is summed in the y direction producing an intensity profile in x for each image in the sequence (**Figure 2**). A detection threshold of the particles is determined as 1 standard deviation of the threshold image pixel intensities above the baseline of 0. A binary particle signal is created where 1 indicates the presence of a particle and 0 indicates no particle.

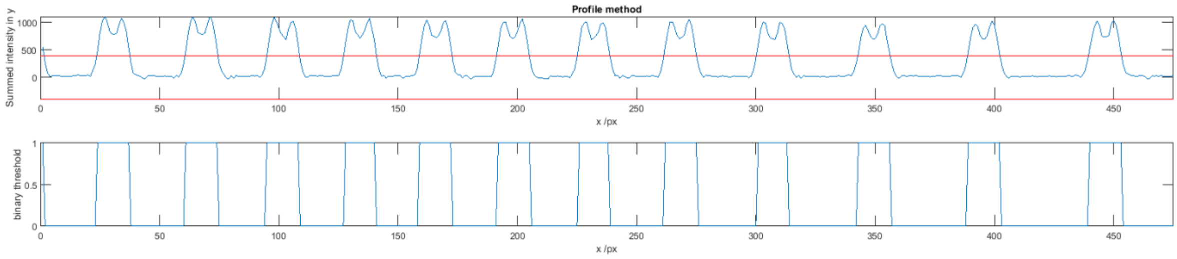


*Figure 2: Pixel intensity summed in y and plotted along x for an individual image from a video sequence (top). Binary particle signal with 1 indicating presence of a particle and 0 indicating no particles (bottom).*

The pitch between binary particle signal peaks is used to calculate the mean particle pitch and standard deviation. The total number of particle signal peaks in a given video sequence is used to calculate frequency.

A second method is used to check that the majority of particles detected are from circular objects. The MATLAB function 'imfindcircles' is used to detect circular shapes and record their position and diameter (**Figure 3**).

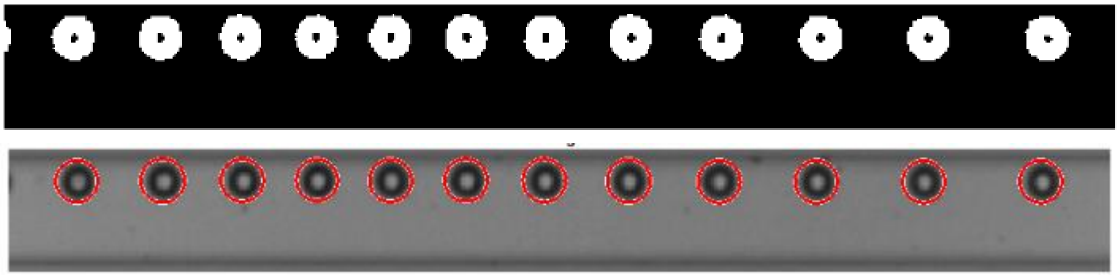

*Figure 3: Black and white image of the particles in the microchannel (top). Circular shapes detected with the MATLAB function 'imfindcircles' (bottom).*

The code outputs are saved as .xlsx files, with the following prefixes followed by the image sequence name:

*binary_particle_signal_intprof_filename.tif.xlsx*

This can be used to plot the displacement resolved binary particle signal for all frames (**Figure 4**).

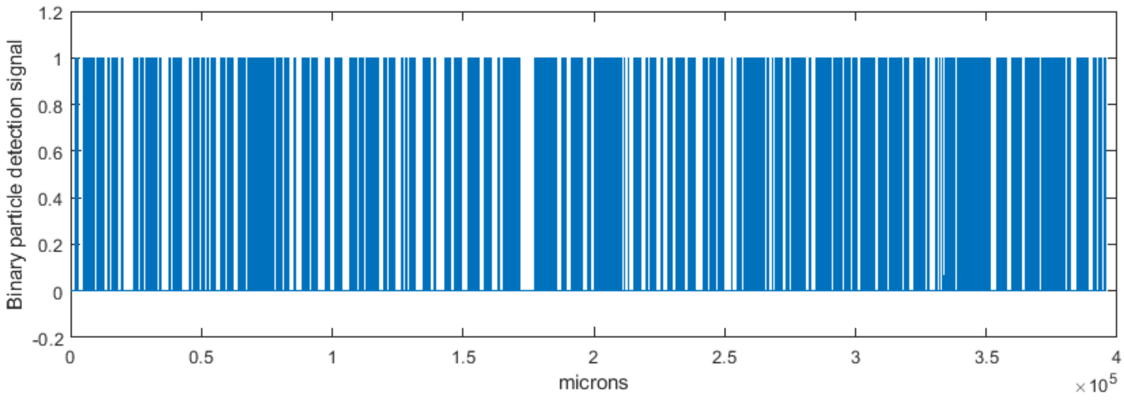


*Figure 4: Displacement resolved binary particle signal ($10^5$ microns = 1 second).*

*Spacing_intprof_filename.tif.xlsx*

The collected particle pitch measurements can be plotted as a histogram (**Figure 5**).

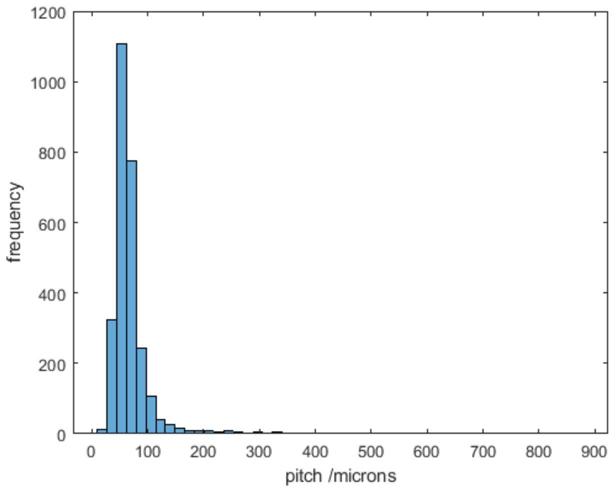


*Figure 5: Frequency distribution of particle spacing.*

*Statistics_intprof.tif.xlsx*

The key statistics for the image sequence and comparison of the intensity profile and circle detection methods.

| Measure | Intensity Profile Method | Circle Method |
|---|---|---|
| count | 3091 | 3311 |
| count rate (/s) | 1236.0 | 1324.0 |
| Spacing mean (px) | 37.7 | 35.4 |
| Spacing sd (px) | 30.5 | 30.2 |
| spacing mean (um) | 73.1 | 68.7 |
| Spacing sd (um) | 59.1 | 58.6 |

*Table 1: .xlsx code output showing the key statistics.*

#### MATLAB CODE

```
clear all; close all;clc;

%PREPARE IMAGE IN IMAGE J
%1)Crop image to just straight channel region.
%2)Rotate image to horizontal flowing right to left
%3)Save as greyscale tif stack in same folder as Drop_frequency_vx.m file
%%%%%%%%%%%%%%%%%%%%%%%%%%%%%%%%%%%%%%%%%%%%%%%%%%%%%%%%%%%%%%%%%%%%%%%%%%%%%%
%Input in MATLAB
%%%%%%%%%%%%%%%%%%%%%%%%%%%%%%%%%%%%%%%%%%%%%%%%%%%%%%%%%%%%%%%%%%%%%%%%%%%%%%
%1)Input full file name between the '' quotation markers
data='6min_6rpm_30um_192x480_1200fps.tif'; %e.g '1minhorizontalmag_crop.tif'

info = imfinfo(data);   %looks up image file data
num_images = numel(info); %number of images in tif stack
%num_images = 500;

%For each variable input the number to the right of the equals sign e.g image_of_interest=4;
image_of_interest=63;      %2)Input image number for display as example image
thresh_image=63;           %3)Input image number with multiple beads in to use to calculate
the threshold
background_image=1;        %4)Input image number which has no beads in to use for background
flattening of image illumination
dec=7;                      %5)Input number of frames to take samples at
frame_rate=1200 ;            %6)Input frame rate of high speed video in fps
scale=0.5154 ;               %7)Input image scale calibration in px/um
min_bead_diam=7;           %9)Input the minimum expected bead diameter in px, used as a
threshold to exclude multipeak abberations
max_bead_diam=30;           %10)Input the maximum expected bead diameter in px

DISPLAY_IMAGE_FOR_CIRCLE_THRESHOLD=1 ;      %11)Input 1 to activate figure for circle method
thresholding selection or 0 not to
binary_threshold_level=8 ; %12) Input the pixel intensity threshold level between 0-256 used
to convert greyscale images to black and white

EXPORT_FILES=1 ;            %13) Input 1 to export xls files or input 0 to not export

%14)PRESS F5 OR CLICK ON RUN AT THE TOP BAR
%The data generated from the code will be displayed in the command window
%The key dat a will also be saved in xls files
%%%%%%%%%%%%%%%%%%%%%%%%%%%%%%%%%%%%%%%%%%%%%%%%%%%%%%%%%%%%%%%%%%%%%%%%%%%%%%
%CODE STARTS
%%%%%%%%%%%%%%%%%%%%%%%%%%%%%%%%%%%%%%%%%%%%%%%%%%%%%%%%%%%%%%%%%%%%%%%%%%%%%%
%% Read images and background subtract
tic;
for i=1:num_images %Reads all images in file
    I(:,:,i)=imread(data,i,'info',info);
end
time1_imread=toc;

I_bs=((imread(data,background_image,'info',info))-I); %subtract background from each image

I_bs_ymed(:,:)=double(median(I_bs)); %median of y direction for each image
I_bs_ysum_med=[I_bs_ymed*size(I,1)]';%summed median by pixel number in y direction
%% Circle detection method
if DISPLAY_IMAGE_FOR_CIRCLE_THRESHOLD==1 %displays a bg subtracted image and histogram used to
find circle method detection threshold
figure(99)
subplot(2,1,1);
imshow(I_bs(:,:,image_of_interest));
title(sprintf('Background subtracted image of interest %d',image_of_interest));

subplot(2,1,2);
imhist(I_bs(:,:,image_of_interest)); % displays histogram of background subtracted image
intensities
xlabel('pixel intensity');
ylabel('frequency');
title('Find local minima in pixelintensity between histogram peaks for background and bead')
else
end

binary_threshold_level=binary_threshold_level/256; %threshold for image binarisation of 8bit
images on scale 1-0
radiusRange=[floor(min_bead_diam/2) floor(max_bead_diam/2)]; %radius min and max for circle
detection method
```

```
circle_gap_vec=[];
for i=1:num_images
BW(:,:,i) = imbinarize(I_bs(:,:,i),binary_threshold_level); %converts background subtracted
image to binary black and white
[centers,radii] = imfindcircles(BW(:,:,i),radiusRange); %finds circular shapes in binary image
if size(centers,1)>0
    centers=[centers radii]; %sorts bead coordinates and raddii by ascending x position
    centers=sortrows(centers,1);%puts x coordinates in ascending order
    radii=centers(:,3);
    centers(:,3)=[];
    centers_rounded=round(centers);
    gap=diff(centers(:,1)); %finds gap between x coordinates
    circle(i).gap=gap;
    circle_gap_vec=[circle_gap_vec;gap]; %creates continuous vector of gap measurements
    circle(i).centers_rounded=centers_rounded; %x coordinates rounded to nearest pixel
else
end
circle(i).number=size(centers,1);
circle(i).centers=centers;
circle(i).radii=radii;
end

gap_image=[];
for i=1:dec:num_images
    if circle(i).number>1;
    gap_image=[gap_image i];
else
    end
end

circle_gap_vec_dec=[];
circle_centre_xpos=[];
radii_vec_dec=[];
m=0;
for i=1:dec:num_images;
    m=m+1;
    circle_gap_vec_dec=[circle_gap_vec_dec; circle(i).gap(:)]; %puts all measured gaps into
continuous vector
    radii_vec_dec=[radii_vec_dec; circle(i).radii(:)]; %puts all radii measurements into
continuous vector
    if size(circle(i).centers,1)>0; %puts all bead x positions into continuous vector
        circle_centre_xpos=[circle_centre_xpos; circle(i).centers_rounded(:,1)+size(I,2)*(m-
1)];
    else
    end
end
radii_vec_dec=round(radii_vec_dec);%rounds radii vector to nearest pixel
circle_gap_vec_dec_um=circle_gap_vec_dec./scale; %gap vector in microns
circle_binary_signal=zeros(ceil(num_images/dec)*size(I,2),1);
circle_binary_signal(circle_centre_xpos)=1; % Creates binary signal with detected bead
position indicated by a 1

for i=1:size(circle_centre_xpos,1); % Uses radii to plot the width of the bead signal in the
binary signal.
        circle_binary_signal(circle_centre_xpos(i)-
radii_vec_dec(i):circle_centre_xpos(i)+radii_vec_dec(i))=1;
end

circle_count_vec=[]; %puts counts for each frame into continuous vector
for i=1:num_images;
    circle_count_vec=[circle_count_vec; circle(i).number(:)];
end

%Circle method statistics
circle_gap_av=mean(circle_gap_vec_dec); %mean of all measured spacings
circle_gap_av_um=circle_gap_av/scale; %mean of all measured spacings in microns
circle_gap_sd=std(circle_gap_vec_dec); %spacing standard deviation in pixels
circle_gap_sd_um=circle_gap_sd/scale; %spacing standard deviation in microns
circle_count=sum(circle_count_vec(1:dec:num_images)); % bead count
circle_count_rate=circle_count/(num_images/frame_rate); %bead count rate

%% Intensity profile method
ys(:,:)=sum(I_bs,1); %takes sum of px intensity in y direction for each image in sequence
producing a summed intensity profile in x for eachi mage
ys=ys';
ys=ys-I_bs_ysum_med; %background subtraction of each intensity profile.
```

```
% by=sum(imread(data,background_image,'info',info)); %reads background image for a background
illumination profile
ys_av=mean(ys,2); % average value for each image profile
ys_sd=std(ys,0,2); % SD for each image profile
thresh=1*ys_sd(thresh_image); % Calculates threshold as 1 x SD from baseline of 0

for i=1:num_images %converts all x profiles to a binary profile
    ybin(ys(i,:)>-thresh & ys(i,:)<thresh)=0; %x coordinates within threshold
    ybin(ys(i,:)<=-thresh | ys(i,:)>=thresh)=1; %x coordinates exceeding threshold
    K(i,:)=ybin; % new matrix of binary x profiles
end

y_col=ys';%Raw signal matrix in column
K_col=K'; %Binary signal matrix in column
y_col_dec=y_col(:,1:dec:size(y_col,2)); %Decimated raw signal matrix
K_col_dec=K_col(:,1:dec:size(K_col,2)); %Decimated binary signal matrix
y_vec=y_col(:); %Raw intensity profile not decimated contin vector
y_vec_dec=y_col_dec(:); %Raw intensity profile decimated contin vector
K_vec=K_col(:); % Binary bead detection signal continuous vector not decimated
K_vec_dec=K_col_dec(:); %Binary bead detection signal continuous vector decimated
K_diff=diff(K,1,2); %Differential, positive slope =1 negative slope =-1

for i=1:num_images
K_a=find(K_diff(i,:)==1)+1; %Peak start position
K_b=find(K_diff(i,:)==-1);  %Peak finish position
K_a_d=diff(K_a);            %Difference between peak starts
K_a1=K_a(find(K_a_d>min_bead_diam)+1); %Finds position of peak starts further than min bead
diameter
K_a2=K_a(find(K_a_d<min_bead_diam)+1);  %Finds positon of peaks closer than min bead diameter
id=find(K_a_d<min_bead_diam)+1; %Finds position of internal peaks
K_a3=K_a;
K_a3(id)=[]; % Removes internal peaks from K_a
Spacing(i)=mean(diff(K_a3)); % Mean spacing between peak starts
Spacing_sd(i)=std(K_a3);
Number_beads(i)=size(K_a3,2);
intprof(i).gap=diff(K_a3);
end

intprof_gap_vec_dec=[];
m=0;
for i=1:dec:num_images;
    m=m+1;
    intprof_gap_vec_dec=[intprof_gap_vec_dec; intprof(i).gap(:)]; %puts all measured gaps into
continuous vector
end

%Intesity profile method statistics
intprof_gap_av=mean(intprof_gap_vec_dec);
intprof_gap_sd=std(intprof_gap_vec_dec);
intprof_gap_av_um=mean(intprof_gap_vec_dec)/scale;
intprof_gap_sd_um=std(intprof_gap_vec_dec)/scale;
Num_spaces_sample=Number_beads(1:dec:num_images)-1; %Total number of spaces between beads
measured. -1 removes measurements from lone beads
Total_num_spaces=sum(Num_spaces_sample(find(Number_beads(1:dec:num_images)>0)));
intprof_count=sum(Number_beads(1:dec:num_images)) ;%Bead count
intprof_count_rate=intprof_count/(num_images/frame_rate);
%% Background subtraction and thresholding figures
%image_of_interest=4001;
figure(1)
subplot(5,1,1)
J=imread(data,image_of_interest,'info',info);
imshow(J);
title(sprintf('Circle method.Image of interest %d',image_of_interest));
%viscircles(circle(image_of_interest).centers(:,1:2),circle(image_of_interest).radii);

subplot(5,1,2)
imshow(I_bs(:,:,image_of_interest));

subplot(5,1,3)
imshow(BW(:,:,image_of_interest));

subplot(5,1,4)
plot(ys(image_of_interest,:));
hold on
threshold_line=refline([0 thresh]);
threshold_line.Color='r';
threshold_line2=refline([0 -thresh(1)]);
```

```
threshold_line2.Color='r';
hold off
xlim([0 size(J,2)]);
xlabel('x /px');
ylabel('Summed intensity in y');
title('Profile method');

subplot(5,1,5)
plot(K(image_of_interest,:));
xlim([0 size(J,2)]);
xlabel('x /px');
ylabel('binary threshold');

if DISPLAY_IMAGE_FOR_CIRCLE_THRESHOLD==1
m=0;
for i=1:50:numel(gap_image)
m=m+1;
    figure_number=10+m;
figure(figure_number);
subplot(5,1,1)
J=imread(data,gap_image(i),'info',info);
imshow(J);
title(sprintf('Circle method.Image of interest %d',gap_image(i)));
viscircles(circle(gap_image(i)).centers(:,1:2),circle(gap_image(i)).radii);

subplot(5,1,2)
imshow(I_bs(:,:,gap_image(i)));

subplot(5,1,3)
imshow(BW(:,:,gap_image(i)));

subplot(5,1,4)
plot(ys(gap_image(i),:));
hold on
threshold_line=refline([0 thresh]);
threshold_line.Color='r';
threshold_line2=refline([0 -thresh(1)]);
threshold_line2.Color='r';
hold off
xlim([0 size(J,2)]);
xlabel('x /px');
ylabel('Summed intensity in y');
title('Profile method');

subplot(5,1,5)
plot(K(gap_image(i),:));
xlim([0 size(J,2)]);
xlabel('x /px');
ylabel('binary threshold');
end
else
end
%% Statistics figures check
figure(2)
subplot(3,1,1)
plot(1:dec:num_images,Number_beads(1:dec:num_images));
xlabel('Image number');
ylabel('Number of beads/frame');
title('Number of beads per frame');

subplot(3,1,2)
plot(1:dec:num_images,Spacing(1:dec:num_images));
xlabel('image number');
ylabel('Mean bead spacing /px');
title('Average bead spacing for each frame. Data gap indicates no bead or single bead in
frame');

subplot(3,1,3)
histogram(intprof_gap_vec_dec,floor(numel(intprof_gap_vec_dec)/2));
xlabel('spacing /px');
ylabel('frequency');
title('Frequency distribution of bead spacing');
hold on
histogram(circle_gap_vec_dec,floor(numel(intprof_gap_vec_dec)/2));
hold off
legend('Intensity profile method','Circle detect method');
%% Detailed figures
```

```
start_frame=1;
finish_frame=10;
nbins=50;

indices=[(size(I,2):1:size(I,2)+numel(K_vec_dec)-1)/size(I,2)]';

figure(3)
subplot(2,1,1)
plot((1:1:numel(y_vec))/size(I,2),y_vec);
hold on
threshold_line=refline([0 thresh]);
threshold_line.Color='r';
threshold_line2=refline([0 -thresh(1)]);
threshold_line2.Color='r';
hold off
%xlim([1 10000]);
xlabel('Frame number');
ylabel('Binary bead detection signal');
title('Intensity profile method: Raw signal all frames');

subplot(2,1,2)
plot((1:1:numel(K_vec))/size(I,2),K_vec);
ylim([-0.2 1.2]);
%xlim([1 10000]);
xlabel('Frame number');
ylabel('Binary bead detection signal all frames');
title('Intensity profile method: Binary signal all frames');

figure(4)
subplot(2,1,1);
plot(K_vec_dec);
ylim([-0.2 1.2])
%xlim([start_frame finish_frame])
xlabel('Pixels');
ylabel('Binary bead detection signal dec');
title('Intensity profiled detection method');

subplot(2,1,2);
plot(circle_binary_signal);
ylim([-0.2 1.2]);
xlabel('Pixels');
ylabel('Binary bead detection signal');
title('Circle detection method');

figure(5)
plot(K_vec_dec);
ylim([-0.2 1.2]);
%xlim([2000 5000]);
xlabel('Pixels');
ylabel('Binary particle detection signal');

figure(6)
plot((1:1:numel(K_vec_dec))./scale,K_vec_dec);
ylim([-0.2 1.2]);
% xlim([0 num_images/4]);
xlabel('microns');
ylabel('Binary particle detection signal');

figure(7)
plot(indices*7*1000/frame_rate,K_vec_dec);
ylim([-0.2 1.2]);
xlim([0 500]);
ylim([0 1]);
xlabel('Time (ms)');
%ylabel('Binary bead detection signal');
set(figure(7), 'Position', [1, 1, 1000, 200]);
set(gca,'YTickLabel',[],'TickDir','out')
print('barcode','-depsc');

figure(8)
histogram(intprof_gap_vec_dec,nbins);
xlabel('pitch /px');
ylabel('frequency');

figure(9)
histogram(intprof_gap_vec_dec/scale,nbins);
xlabel('pitch /microns');
```

```
ylabel('frequency');

%% Export files
if EXPORT_FILES==1
%intprof export
col_header={'frame' 'Pixels','Microns','binary signal'};
binary_bead_signal_intprof_output=num2cell([indices [1:1:numel(K_vec_dec)]'
[(1:1:numel(K_vec_dec))./scale]' K_vec_dec]);
binary_bead_signal_intprof_output=[col_header; binary_bead_signal_intprof_output];
xlswrite(sprintf('binary_bead_signal_intprof_%s.xlsx',data),[binary_bead_signal_intprof_output
]);

xlswrite(sprintf('Spacing_intprof_%s.xlsx',data),[intprof_gap_vec_dec]);

%statistics export
statistics_output=num2cell([intprof_count circle_count;...
    intprof_count_rate circle_count_rate;...
    intprof_gap_av circle_gap_av;...
    intprof_gap_sd circle_gap_sd;...
    intprof_gap_av_um circle_gap_av_um;...
    intprof_gap_sd_um circle_gap_sd_um]);
col_header={'Intensity profile method','Circle method'};
row_header={'count'; 'count rate (/s)';'Spacing mean (px)';'Spacing sd (px)';'spacing mean
(um)';'Spacing sd (um)';};
statistics_output=[{' '} col_header; row_header statistics_output];

xlswrite(sprintf('Statistics_%s.xlsx',data),[statistics_output]);
else
end

% %circles export
% col_header={'Pixels','Microns','binary signal'};
% binary_bead_signal_circles_output=num2cell([[1:numel(circle_binary_signal)]'
[(1:numel(circle_binary_signal))./scale]' circle_binary_signal]);
% binary_bead_signal_circles_output=[col_header; binary_bead_signal_circles_output];
%
xlswrite(sprintf('binary_bead_signal_circles_%s.xlsx',data),[binary_bead_signal_circles_output
]);
%
% xlswrite(sprintf('Spacing_circles_%s.xlsx',data),[circle_gap_vec_dec]);
```

### Appendix III: Droplet Encapsulation MATLAB Code

#### Code Overview

This code identifies and measures the size of microfluidic droplets in a microfluidic channel and the number of particles and cells or 'cell'-sized beads within these. The original video file is directly imported into MATLAB variable space and the cropping function of MATLAB is used to define the region of interest. It is preferable to synchronise the frame rate of the camera with the droplet generation rate, so that a new droplet will appear at approximately the same location as the previous droplet in the next frame. However, if this cannot be achieved, it is recommended to use a lower camera frame rate. First, the region of interest is analysed to find droplets. When droplets are found the code searches a droplet that is fully contained in the region of interest and saves the geometric centre and the radius. Then the original region of interest is further cropped using this information so that only one droplet is now captured.

Two different detection methods are used to capture cells/small particles whereas large particles can be accurately found by using only one detection method. Cells/small particles can be partially hidden in the droplet interface or out of focus and therefore the use of two separate methods enables accurate particle identification. The first method for cell/small particle detection involves a user input intensity threshold together with the MATLAB function 'imfindcircles' to locate the particle. Then, a second method with a different intensity threshold and the MATLAB function 'regionprops' is used. In the case of detecting the particle using both methods, the code only saves the information once to prevent oversampling. Similarly, large particles are detected using a third intensity threshold and the MATLAB function 'imfindcircles'. Again, if large particles are also detected by the small particle detection method these are removed from the list of detected small particles. Once the droplet is analysed, the cropped image of the droplet is classified and saved in the corresponding folder for visual verification.

An image generated from the code when run under 'PLOT mode' (PLOT=1;) is shown in **Figure 1**. These can be used to check the accuracy of the detection algorithm. When the script is run in this mode it will generate one figure per frame and therefore the amount of frames should be kept low to reduce memory load. For automatic processing the code should be run with PLOT=0.

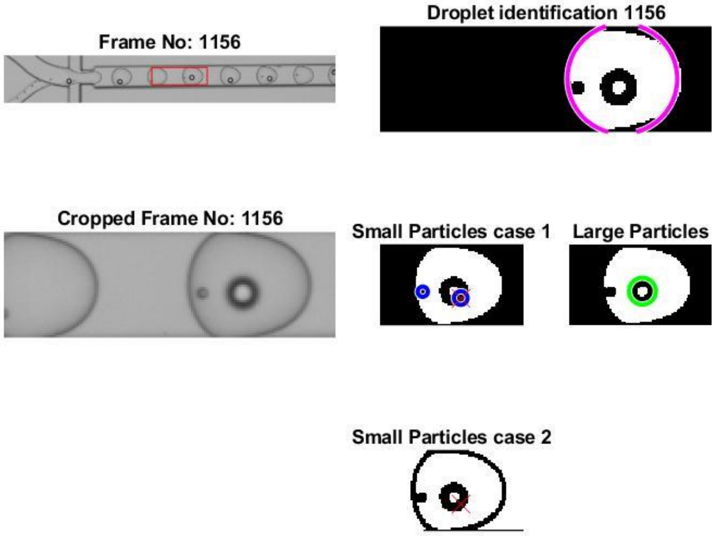


***Figure 1:*** *Image obtained from the script in 'PLOT mode'. Frame 1156 shows the identification of a droplet with a large and small particle. In this instance a large particle is detected using both small and large methods with the red cross indicating particle rejection.*

#### MATLAB CODE

```
clear all; close all; clc;
%Load video
[file,path]=uigetfile('.avi','Select the .AVI file');

namevideo1=strcat([path file]);
f=waitbar(0,'Loading video...');

vid1 = VideoReader(namevideo1);
waitbar(0.5, f,'Loading video...');

vid1 = read(vid1(:,:,:)); %Save the video in Matlab space
waitbar(0.9, f,'Almost there!');

[pixely,pixelx,tt,frames]=size(vid1);
waitbar(1,f,'Done!');
pause(1)
close(f)

%Generate folders
New_Directory = fullfile('C:\Users\luibl\Documents\MATLAB\Cells');  %full path of directory. Good way to do it
foldernew = fullfile(New_Directory, file(1:end-4));
mkdir(foldernew);
folder_empty = fullfile(foldernew, 'Empty');
mkdir(folder_empty);

folder_one_small = fullfile(foldernew, 'One_small');
mkdir(folder_one_small);

folder_one_large = fullfile(foldernew, 'One_large');
mkdir(folder_one_large);

folder_one_both = fullfile(foldernew, 'One_both');
mkdir(folder_one_both);

folder_two_small_one_large = fullfile(foldernew, 'Two_small_one_large');
mkdir(folder_two_small_one_large);

folder_two_large_one_small = fullfile(foldernew, 'Two_large_one_small');
mkdir(folder_two_large_one_small);

folder_two_both = fullfile(foldernew, 'Two_both');
mkdir(folder_two_both);

folder_two_small = fullfile(foldernew, 'Two_small');
mkdir(folder_two_small);

folder_two_large = fullfile(foldernew, 'Two_large');
mkdir(folder_two_large);

folder_three_small = fullfile(foldernew, 'Three_small');
mkdir(folder_three_small);

folder_three_large = fullfile(foldernew, 'Three_large');
mkdir(folder_three_large);
```

```
folder_three_small_one_large = fullfile(foldernew, 'Three_small_one_large');
mkdir(folder_three_small_one_large);

folder_three_large_one_small = fullfile(foldernew, 'Three_large_one_small');
mkdir(folder_three_large_one_small);


%%
close all
%Droplet diameter
Dmin=10;
Dmax=50;
%Small droplet detection using findcircle (for further parameters go to
%line 92
dmin=1;
dmax=4;
%Small droplet detection using regionprops (for further parameters go to
%line 128
Amin=2;
Amax=70;
%Large droplet detection sing findcircle (for further parameters go to
%line 174
dlmin=5;
dlmax=10;
%Thresholds for detection
Threshold2=90;
Threshold3=85;

Cent_tolerance=10; %tolerance to identify the duplicated droplets

rectL=[400,33,150,47]; %Proposed Region of Interest [x1,y1,Deltax,Deltay]
PLOT=0; % PlOT=1 to get figures from every snapshot;
%If PLOT=1 and the number of frames is too large your computer will crash!
FrameIni=1; %First frame to check
FrameEnd=5000; %Last frame to check

%Initialize variables
RR=[];
cc=0;
cim=0;
Dropletsfound=[];

Case_Empty=0;
Case_One_Small=0;
Case_One_Large=0;
Case_One_Both=0;
Case_Two_Small_One_Large=0;
Case_Two_Large_One_Small=0;
Case_Two_Both=0;
Case_Two_Small=0;
Case_Two_Large=0;
Case_Three_Large=0;
Case_Three_Small=0;
Case_Three_Large_One_Small=0;
Case_Three_Small_One_Large=0;

%Loop from first to last frame
```

```
for i=FrameIni:FrameEnd
  cim=cim+1;
  try
    IOri=vid1(:,:,1,i); %Load image

    Iback= imcrop(IOri,rectL); %Crop image
    Istart=Iback;
    %manual threhold
    % threshold
    t = 145; %Modify this value according to setup
    % find values below
    ind_below = (Iback < t);
    % find values above
    ind_above = (Iback >= t);
    % set values below to black
    Iback(ind_below) = 0;
    % set values above to white
    Iback(ind_above) = 255;

    inverseBW = uint8(255) - Iback;

    BW=inverseBW;

    BW2 = imfill(BW,'holes');
    BW_interface=imfill(BW2-BW, 'holes');

    Icropv2=Iback.*uint8((BW_interface));

    %Find droplet
    [centL,radL]=imfindcircles(Icropv2,[Dmin
Dmax],'ObjectPolarity','bright','Sensitivity',0.9,'EdgeThreshold',0.1,'method', 'TwoStage'); %Find droplet

    if isempty(radL)==0 %if Region Of Interest is not empty
      ToOri=centL(:,1)-radL;
      ToEnd=centL(:,1)+radL;
      chec=ismember(find(ToOri>0), find(ToEnd< rectL(3)));
      radLfull=[];
      centLfull=[];
      flag=0;

      for ki=1:length(chec)
        if chec(ki)==true
          radLfull=[radLfull; radL(ki)];
          centLfull=[centLfull; centL(ki,:)];
        else
          flag=flag+1;
        end
      end
      %Choose the first droplet to appear (to avoid counting them twice)
      [~,idx] = sort(centLfull(:,1)); % sort just the first column
      sortedcentL = centLfull(idx,:);   % sort the whole matrix using the sort indices
      sortedradL = radLfull(idx,:);

      if  ismember(1, chec)%Any droplet is not empty

        if PLOT==1
          figure
```

```
        subplot(3,4,1:2); %Original image with ROI section
        imshow(IOri)
        hold on
        rectangle('Position',rectL,'EdgeColor', 'r')
        title(['Frame No:' ' ' num2str(i)]);

        subplot(3,4,5:6); %Original image with ROI section
        imshow(Istart)
        hold on
        rectangle('Position',rectL,'EdgeColor', 'r')
        title(['Cropped Frame No:' ' ' num2str(i)]);

        subplot(3,4,3:4);% ROI section with droplet detected
        imshow(Icropv2)
        hold on
        viscircles(centLfull,radLfull,'Color','m');
        title(['Droplet identification' ' ' num2str(i)]);
    else
    end
    RR=[RR; sortedradL(1)];
    Dropletsfound=[Dropletsfound; 1];

    %Crop image according to first droplet found
    Iback = Iback(:,max(1,sortedcentL(1)-radL(1)-20):min(rectL(3), sortedcentL(1)+radL(1)+20));
    Icropv2 = Icropv2(:,max(1,sortedcentL(1)-radL(1)-20):min(rectL(3), sortedcentL(1)+radL(1)+20));
    Istartv2 = Istart(:,max(1,sortedcentL(1)-radL(1)-20):min(rectL(3), sortedcentL(1)+radL(1)+20));

    %Find small particles
    Ibackwhite=Iback;
    [cent,rad]=imfindcircles(Icropv2,[dmin
dmax],'ObjectPolarity','dark','Sensitivity',0.75,'EdgeThreshold',0.075,'method', 'TwoStage');
    centS1=cent;
    radS1=rad;
    if PLOT==1
        subplot(3,4,7); %Small particles with findcircles
        imshow(Icropv2);
        viscircles(cent,rad,'Color','b');
        title(['Small Particles case 1'])
        hold on
    else
    end

    cc=0;
    %Save the data found
    if isempty(rad)==0
        for j=1:length(rad)
            cc=cc+1;
            regFsub8(cim).n(cc).Rad=rad(j);
            regFsub8(cim).n(cc).Centroid=cent(j,:);
        end
    else
        regFsub8(cim).n=[];
    end

    Centroids8=[0 0];
    for k=1:length(regFsub8(cim).n)
        Centroids8=[Centroids8; regFsub8(cim).n(k).Centroid];
```

```
            end

            IallWhite2=Ibackwhite;
            IallWhite2(IallWhite2>Threshold2)=300; %Manual threshold
            IallWhite2=imgaussfilt(IallWhite2,'FilterSize',1); %Gaussian filter

           %These lines to remove small particles (noise)
            BW = imbinarize(IallWhite2);
            BW= (BW-1)*-1;
            CC = bwconncomp(BW,4);
            S = regionprops(CC,'Area','Eccentricity','centroid');
            L = labelmatrix(CC);
            BWNew = ismember(L, find([S.Area] <= Amax & [S.Area] >= Amin));%;AxisLength','Eccentricity','Area');
            regBsub11 =
regionprops(BWNew,'centroid','MajorAxisLength','MinorAxisLength','Eccentricity','Area');
            if PLOT==1
               subplot(3,4,11) %Small particles with regprops
               imshow(IallWhite2)
               title(['Small Particles case 2'])
               hold on;
            else
            end
            cc=0;
            %Evaluation of coincidence between cases to avoid counting
            %particles twice
            if isempty (regBsub11)==0
               for j=1:length(regBsub11)
                  [mv1,mv2]=size(Icropv2);
                  if regBsub11(j).MajorAxisLength>2 && regBsub11(j).Area<100 &&
regBsub11(j).Centroid(1)<(mv2-5) && regBsub11(j).Centroid(1)>2 && regBsub11(j).Centroid(2)>2 &&
regBsub11(j).Centroid(2)<mv1-5

                     XX=regBsub11(j).Centroid(1)-Centroids8(:,1);
                     YY=regBsub11(j).Centroid(2)-Centroids8(:,2);

                     if min(sqrt(XX.^2+YY.^2))>5
                        cc=cc+1;
                        regFsub11(cim).n(cc)=regBsub11(j);
                        Cent(cc,:)=regBsub11(j).Centroid;
                        if PLOT==1
                           plot(Cent(cc,1),Cent(cc,2),'ro');
                        else
                        end
                     else
                     end


                  else

                  end
               end
            else
            end

            if cc==0
               regFsub11(cim).n=[];
            else
```

```
            end

            IallWhite=Ibackwhite;
            IallWhite(IallWhite>Threshold3)=300;
            [cent,rad]=imfindcircles(Icropv2,[dlmin
dlmax],'ObjectPolarity','dark','Sensitivity',0.9,'EdgeThreshold',0.5,'method', 'TwoStage'); %Find Large Bead
            [m,n]=size(cent);

            if PLOT==1
               subplot(3,4,8) %Large particles with findcircles
               imshow(Icropv2);
               hold on
               viscircles(cent,rad,'Color','g');
               title(['Large Particles'])
            else
            end

            %If we find a large particle we verify that this was not
            %identified as a small particle previously. If that is the
            %case we delet the 'small' particle.

            if isempty(rad)==0
               for j=1:m
                  XX=cent(j,1)-Centroids8(:,1);
                  YY=cent(j,2)-Centroids8(:,2);
                  [val2,loc2]=min(sqrt(XX.^2+YY.^2));
                  VAL=sqrt(XX.^2+YY.^2);
                  SmallVAL=find(VAL<Cent_tolerance);
                  regFsub9(cim).n(j).Centroid=cent(j,:);
                  regFsub9(cim).n(j).Rad=rad(j);

                  if PLOT==1 && length(SmallVAL)>=1
                     subplot(3,4,11)
                     hold on

                     for ko=1:length(SmallVAL)
                        plot(Centroids8(SmallVAL(ko),1),Centroids8(SmallVAL(ko),2),'rx','MarkerSize',12)
                        regFsub8(cim).n(SmallVAL(ko)-1).Centroid=[];
                        regFsub8(cim).n(SmallVAL(ko)-1).Rad=[];
                        regFsub8(cim).n(SmallVAL(ko)-1)=[];
                     end

                     subplot(3,4,7)
                     hold on
                     for ko=1:length(SmallVAL)
                        plot(Centroids8(SmallVAL(ko),1),Centroids8(SmallVAL(ko),2),'rx','MarkerSize',12)
                     end
                  else
                  end

               end
            else
               regFsub9(cim).n=[];
               Dropletsfound=[Dropletsfound; 0];
            end
         else
         end
```

```
if PLOT==0
    SmallThr1=regFsub8; %Small Beads found with detection 1
    SmallThr2=regFsub11;%Small Beads found with detection 2
    LargeThr3=regFsub9;%Large Beads found with detection 3
    DropRad=RR; %%Size of droplet


    %Save droplet image in corresponding folder
    if (length(SmallThr1(cim).n)+length(SmallThr2(cim).n)==0 && length(LargeThr3(cim).n)==0)
        Case_Empty=[Case_Empty +1];
        fullFileName = fullfile(folder_empty, ['Empty_' num2str(length(Case_Empty)) '.png']);
        imwrite(Istartv2, fullFileName);
    elseif (length(SmallThr1(cim).n)+length(SmallThr2(cim).n)==1 && length(LargeThr3(cim).n)==0)
        Case_One_Small=[Case_One_Small +1];
        fullFileName = fullfile(folder_one_small, ['One_small' num2str(length(Case_One_Small)) '.png']);
        imwrite(Istartv2, fullFileName);
    elseif (length(SmallThr1(cim).n)+length(SmallThr2(cim).n)==0 && length(LargeThr3(cim).n)==1)
        Case_One_Large=[Case_One_Large +1];
        fullFileName = fullfile(folder_one_large, ['One_large' num2str(length(Case_One_Large)) '.png']);
        imwrite(Istartv2, fullFileName);
    elseif (length(SmallThr1(cim).n)+length(SmallThr2(cim).n)==1 && length(LargeThr3(cim).n)==1)
        Case_One_Both=[Case_One_Both +1];
        fullFileName = fullfile(folder_one_both, ['One_both' num2str(length(Case_One_Both)) '.png']);
        imwrite(Istartv2, fullFileName);
    elseif (length(SmallThr1(cim).n)+length(SmallThr2(cim).n)==2 && length(LargeThr3(cim).n)==1)
        Case_Two_Small_One_Large=[Case_Two_Small_One_Large +1];
        fullFileName = fullfile(folder_two_small_one_large, ['Two_small_one_large'
num2str(length(Case_Two_Small_One_Large)) '.png']);
        imwrite(Istartv2, fullFileName);
    elseif (length(SmallThr1(cim).n)+length(SmallThr2(cim).n)==1 && length(LargeThr3(cim).n)==2)
        Case_Two_Large_One_Small=[Case_Two_Large_One_Small +1];
        fullFileName = fullfile(folder_two_large_one_small, ['Two_large_one_small'
num2str(length(Case_Two_large_One_small)) '.png']);
        imwrite(Istartv2, fullFileName);
    elseif (length(SmallThr1(cim).n)+length(SmallThr2(cim).n)==2 && length(LargeThr3(cim).n)==2)
        Case_Two_Both=[Case_Two_Both +1];
        fullFileName = fullfile(folder_two_both, ['Two_both' num2str(length(Case_Two_both)) '.png']);
        imwrite(Istartv2, fullFileName);
    elseif (length(SmallThr1(cim).n)+length(SmallThr2(cim).n)==2 && length(LargeThr3(cim).n)==0)
        Case_Two_Small=[Case_Two_Small +1];
        fullFileName = fullfile(folder_two_small, ['Two_small' num2str(length(Case_Two_small)) '.png']);
        imwrite(Istartv2, fullFileName);
    elseif (length(SmallThr1(cim).n)+length(SmallThr2(cim).n)==0 && length(LargeThr3(cim).n)==2)
        Case_Two_Large=[Case_Two_Large +1];
        fullFileName = fullfile(folder_two_large, ['Two_large' num2str(length(Case_Two_Large)) '.png']);
        imwrite(Istartv2, fullFileName);
    elseif (length(SmallThr1(cim).n)+length(SmallThr2(cim).n)==0 && length(LargeThr3(cim).n)>=3)
        Case_Three_Large=[Case_Three_Large +1];
        fullFileName = fullfile(folder_three_large, ['Three_large' num2str(length(Case_Three_Large))
'.png']);
        imwrite(Istartv2, fullFileName);
    elseif (length(SmallThr1(cim).n)+length(SmallThr2(cim).n)>=3 && length(LargeThr3(cim).n)==0)
        Case_Three_Small=[Case_Three_Small +1];
        fullFileName = fullfile(folder_three_small, ['Three_small' num2str(length(Case_Three_Small))
'.png']);
        imwrite(Istartv2, fullFileName);
    elseif (length(SmallThr1(cim).n)+length(SmallThr2(cim).n)==1 && length(LargeThr3(cim).n)>=3)
```

```
                Case_Three_Large_One_Small=[Case_Three_Large_One_Small +1];
                fullFileName = fullfile(folder_three_large_one_small, ['Three_large_one_small'
num2str(length(Case_Three_Large_One_Small)) '.png']);
                imwrite(Istartv2, fullFileName);
            elseif (length(SmallThr1(cim).n)+length(SmallThr2(cim).n)>=3 && length(LargeThr3(cim).n)==1)
                Case_Three_Small_One_Large=[Case_Three_Small_One_Large +1];
                fullFileName = fullfile(folder_three_small_one_large, ['Three_small_one_large'
num2str(length(Case_Three_Small_One_Large)) '.png']);
                imwrite(Istartv2, fullFileName);
            else

            end
        else
        end
    else
    end
  catch
    fprintf(['ERROR IN FRAME:' num2str(i) '\n'])

  end

  if PLOT==0 && i==1
    f=waitbar(0,'Processing video...');

  elseif  PLOT==0 && i<=FrameEnd-1
    waitbar(i/(FrameEnd-FrameIni), f,'Processing video...');

  elseif PLOT==0 && i==FrameEnd
    waitbar(1, f,'Done!');
    pause(0.5)
    close(f)
  end
end

SmallThr1=regFsub8; %Small Beads found with detection 1
SmallThr2=regFsub11;%Small Beads found with detection 2
LargeThr3=regFsub9;%Large Beads found with detection 3
DropRad=RR; %%Size of droplet

%Write summary text file
Mean_DropRad=mean(DropRad);
Std_DropRad=std(DropRad);
Case_Empty=[];
Case_One_Small=[];
Case_One_Large=[];
Case_One_Both=[];
Case_Two_Small_One_Large=[];
Case_Two_Large_One_Small=[];
Case_Two_Both=[];
Case_Two_Small=[];
Case_Two_Large=[];
Case_Three_Large=[];
Case_Three_Small=[];
Case_Three_Large_One_Small=[];
Case_Three_Small_One_Large=[];
for ki=1:length(DropRad)
  if (length(SmallThr1(ki).n)+length(SmallThr2(ki).n)==0 && length(LargeThr3(ki).n)==0)
```

```
        Case_Empty=[Case_Empty ki];
    elseif (length(SmallThr1(ki).n)+length(SmallThr2(ki).n)==1 && length(LargeThr3(ki).n)==0)
        Case_One_Small=[Case_One_Small ki];
    elseif (length(SmallThr1(ki).n)+length(SmallThr2(ki).n)==0 && length(LargeThr3(ki).n)==1)
        Case_One_Large=[Case_One_Large ki];
    elseif (length(SmallThr1(ki).n)+length(SmallThr2(ki).n)==1 && length(LargeThr3(ki).n)==1)
        Case_One_Both=[Case_One_Both ki];
    elseif (length(SmallThr1(ki).n)+length(SmallThr2(ki).n)==2 && length(LargeThr3(ki).n)==1)
        Case_Two_Small_One_Large=[Case_Two_Small_One_Large ki];
    elseif (length(SmallThr1(ki).n)+length(SmallThr2(ki).n)==1 && length(LargeThr3(ki).n)==2)
        Case_Two_Large_One_Small=[Case_Two_Large_One_Small ki];
    elseif (length(SmallThr1(ki).n)+length(SmallThr2(ki).n)==2 && length(LargeThr3(ki).n)==2)
        Case_Two_Both=[Case_Two_Both ki];
    elseif (length(SmallThr1(ki).n)+length(SmallThr2(ki).n)==2 && length(LargeThr3(ki).n)==0)
        Case_Two_Small=[Case_Two_Small ki];
    elseif (length(SmallThr1(ki).n)+length(SmallThr2(ki).n)==0 && length(LargeThr3(ki).n)==2)
        Case_Two_Large=[Case_Two_Large ki];
    elseif (length(SmallThr1(ki).n)+length(SmallThr2(ki).n)==0 && length(LargeThr3(ki).n)>=3)
        Case_Three_Large=[Case_Three_Large ki];
    elseif (length(SmallThr1(ki).n)+length(SmallThr2(ki).n)>=3 && length(LargeThr3(ki).n)==0)
        Case_Three_Small=[Case_Three_Small ki];
    elseif (length(SmallThr1(ki).n)+length(SmallThr2(ki).n)==1 && length(LargeThr3(ki).n)>=3)
        Case_Three_Large_One_Small=[Case_Three_Large_One_Small ki];
    elseif (length(SmallThr1(ki).n)+length(SmallThr2(ki).n)>=3 && length(LargeThr3(ki).n)==1)
        Case_Three_Small_One_Large=[Case_Three_Small_One_Large ki];
    else

    end
end


fileID = fopen([file(1:end-4) '.txt'],'w');
fprintf(fileID,'Empty = %f\r\n',length(Case_Empty));
fprintf(fileID,'One Small = %f\r\n',length(Case_One_Small));
fprintf(fileID,'One Large= %f\r\n',length(Case_One_Large));
fprintf(fileID,'One Small One Large = %f\r\n',length(Case_One_Both));
fprintf(fileID,'Two Small = %f\r\n',length(Case_Two_Small));
fprintf(fileID,'Two Large = %f\r\n',length(Case_Two_Large));
fprintf(fileID,'Two Small One Large = %f\r\n',length(Case_Two_Small_One_Large));
fprintf(fileID,'One Small Two Large = %f\r\n',length(Case_Two_Large_One_Small));
fprintf(fileID,'Two Small Two Large = %f\r\n',length(Case_Two_Both));
fprintf(fileID,'Three or more Large = %f\r\n',length(Case_Three_Large));
fprintf(fileID,'Three or more Small = %f\r\n',length(Case_Three_Small));
fprintf(fileID,'Three or more Large One Small = %f\r\n',length(Case_Three_Large_One_Small));
fprintf(fileID,'Three or more Small One Large= %f\r\n',length(Case_Three_Small_One_Large));
fprintf(fileID,'Total No. Droplets= %f\r\n',length(DropRad));

fprintf(fileID,'-------------\r\n');
fprintf(fileID,'Mean Particle rad [px] = %f\r\n', Mean_DropRad);
fprintf(fileID,'Std Particle rad [px] = %f\r\n', Std_DropRad);
fclose(fileID);
```
